# Simulated spaceflight disrupts the immune-gut-brain axis and drives sex-dependent neuroinflammation, axonal injury, and behavioral deficits

**DOI:** 10.64898/2026.04.09.717543

**Authors:** Marissa Burke, Göknur Kara, Morgan Holcomb, Christopher Mason, Sonia Villapol

**Author notes:** Corresponding author: Sonia Villapol, Ph.D. Center for Neuroregeneration, Department of Neurosurgery, Houston Methodist Research Institute, 6670 Bertner Avenue, R11-212, Houston, TX 77030, USA. ORCID 0000-0002-6174-4113.

## Abstract

Simulated spaceflight perturbs multiple organ systems, yet the integrated impact of spaceflight-relevant stressors on the immune-gut-brain axis remains poorly defined. We used a ground-based model combining hindlimb unloading (HU) with low-dose ionizing radiation (IR; 50 or 100cGy) to quantify neuropathology, peripheral immune phenotypes, intestinal barrier integrity, and behavioral performance in male and female C57BL/6 mice. HU and/or IR induced region-selective neurodegenerative changes consistent with axonal injury across the cortex and major white-matter tracts. In the somatosensory cortex, MAP-2+ neurons were reduced and SMI-312-labeled axonal injury increased, lowering the intact-to-dystrophic axonal area ratio. Long-range fiber pathways (corpus callosum, cingulate gyrus, external capsule) showed robust axonal damage accompanied by gliosis, with elevated Iba-1+ microglia and GFAP+ astrocytes most prominent after HU+IR (100cGy). Peripheral immunophenotyping revealed a sustained, sex-dependent innate inflammatory bias, with expanded CD11b+ myeloid cells and increased TNF-α+ myeloid activation after IR and IR+HU, alongside maladaptive T-cell polarization despite largely unchanged total CD8+ and CD4+ frequencies. In parallel, the gut exhibited architectural remodeling and barrier failure, including altered mucin profiles, reduced ZO-1 tight-junction labeling, and increased CD45+ leukocyte infiltration across the jejunum, ileum, and colon. Behavioral assays demonstrated sex-dependent deficits spanning affective, motor, and cognitive domains, including increased anxiety- and depressive-like behaviors, impaired rotarod performance, reduced recognition memory, and less efficient spatial strategies. Overall, these findings identify a sex-dependent immune-gut-brain vulnerability in which combined HU and low-dose IR drive gut barrier breakdown and immune imbalance that coincide with neuroinflammatory axonopathy and measurable neurobehavioral dysfunction.

## 1. INTRODUCTION

Deep-space exploration will require astronauts to remain healthy and cognitively resilient for extended periods while exposed to environmental stressors that exceed those of typical low Earth orbit missions. Two dominant physical hazards of the spaceflight environment are microgravity and ionizing radiation^1,2^, which together promote immune dysfunction, oxidative stress, DNA damage, and neurobehavioral alterations in human and experimental studies^3–6^. These responses have raised concern that long-duration missions may accelerate biological aging processes and increase vulnerability to neurodegenerative trajectories characterized by persistent neuroinflammation and altered gene signatures^7–9^.

Astronauts and spaceflight analog cohorts also report sensorimotor deficits, mood-related symptoms, and impairments in cognitive speed and accuracy^10–15^. Yet, the biological mechanisms that translate spaceflight stressors into neurobehavioral outcomes remain incompletely defined. Emerging evidence suggests that cognitive dysfunction may reflect a systemic inflammatory state rather than an exclusively brain-restricted process^7,8^. Prolonged exposure to microgravity and radiation challenges immunoregulation across multiple organs and may compromise barrier function, allowing inflammatory signals from peripheral compartments to influence the central nervous system (CNS)^16,17^. Although blood-brain barrier (BBB) disruption has been reported after spaceflight exposure^17,18^, whether this loss of integrity enables peripheral inflammatory mediators to enter or reshape the brain parenchymal environment, and thereby contribute to neurobehavioral impairment, remains insufficiently addressed.

Spaceflight and analog studies consistently demonstrate immune remodeling, including sustained inflammatory phenotypes, altered leukocyte function, and features consistent with immune activation^19–21^. Peripheral inflammation is also strongly linked to neurodegenerative risk^9^. Communication between the periphery and the CNS can occur through humoral pathways and neural routes, including vagal signaling, even in the absence of overt BBB leakage^22^. Moreover, immune cells can traffic between peripheral tissues and the nervous system, notably, gut- or adipose-derived T cell populations have been reported to influence CNS-adjacent immune tone and cytokine production, including interferon-γ and interleukin-17, with potential relevance to behavior and neuroinflammatory disease states^23^. Importantly, sex differences in immune and neurobehavioral outcomes have been reported across spaceflight datasets and models^20,24,25^. Together, these observations suggest that the peripheral immune compartment may be a key driver of CNS inflammation and behavioral change under spaceflight-relevant stress.

A major route through which systemic immune signals may be integrated with neural function is the gut-brain axis (GBA). The vagus nerve provides bidirectional communication between the CNS and the gastrointestinal tract, coordinating parasympathetic tone and interacting with immune and neuroendocrine networks^26,27^. The GBA encompasses gut-associated lymphoid tissue (GALT), the microbiome, intestinal barrier integrity, and microbial and host-derived metabolites and neurotransmitters that shape both peripheral immunity and brain function^28–31^. On Earth, GBA disruption, through intestinal inflammation, dysbiosis, and barrier breakdown, has been linked to neuroinflammation and behavioral dysfunction, and can also compromise BBB integrity^32–37^. Spaceflight studies likewise report altered gut microbial communities, intestinal barrier perturbations, and neuroendocrine dysregulation^38–41^. However, despite these converging lines of evidence, integrated studies that simultaneously profile immune, gut, and brain inflammatory changes alongside behavior following combined microgravity and radiation exposures remain limited.

Here, we test the hypothesis that combined simulated microgravity and low-dose ionizing radiation initiate a sex-dependent immune-gut-brain cascade in which early immune remodeling and intestinal dysfunction precede or accompany neuroinflammation and behavioral function. To address this, we used a terrestrial spaceflight murine model combining hindlimb unloading (HU) to model microgravity-associated mechanical unloading and cephalad fluid shift with low linear energy transfer (LET) X-ray irradiation as a practical proxy for space radiation exposure^2,42^. Because biological effectiveness depends on radiation quality, we selected doses informed by mission-relevant effective dose estimates and by differences between absorbed dose and LET-weighted risk^1,43,44^, while acknowledging that low-LET X-rays do not fully recapitulate high-LET HZE components of galactic cosmic rays (GCR). Using this model, we systematically quantify immune changes, gut pathology and barrier integrity, neuroinflammation and axonal injury, and behavioral outcomes to define a mechanistic immune-gut-brain pathway linking spaceflight stressors to neurobehavioral dysfunction.

## 2. METHODS

### 2.1. Animals

Adult C57BL/6J mice (21 weeks old; male and female; The Jackson Laboratory, Bar Harbor, ME, USA) were housed in the Houston Methodist Research Institute animal facility under a 12 h light/dark cycle with *ad libitum* access to food and water. To minimize the potential radioprotective effects of phytoestrogens^45^, mice were maintained on a phytoestrogen-reduced Teklad 14 diet. All procedures were approved by the Houston Methodist Research Institute Institutional Animal Care and Use Committee (IACUC, Protocol number: IS00007598) and conducted in accordance with institutional guidelines.

### 2.2. Combined hindlimb unloading and low-dose ionizing radiation model

Mice were transferred to custom-built cages and acclimated for 7 days prior to experimental manipulation. HU or sham procedures were performed as previously described^42^, for tail-suspension paradigms, using tail taping and suspension hardware to maintain unloading while allowing access to food and water. Animals were monitored daily throughout the unloading period. HU was maintained for 14 consecutive days. On day 15, mice received whole-body X-ray irradiation (IR) at 0, 50, or 100cGy using an RS-2000 irradiator, after which hindlimbs were reloaded. Blood and behavioral testing were conducted during the subsequent 2-week post-reloading period (**Fig. 1**). X-ray absorbed doses were selected to approximate space-relevant effective dose estimates for galactic cosmic radiation exposure (e.g., ∼15cGy lunar and ∼50cGy Martian), and were implemented as 50 and 100cGy X-ray doses, respectively, based on effective dose considerations^1,46^.

**Figure 1.**
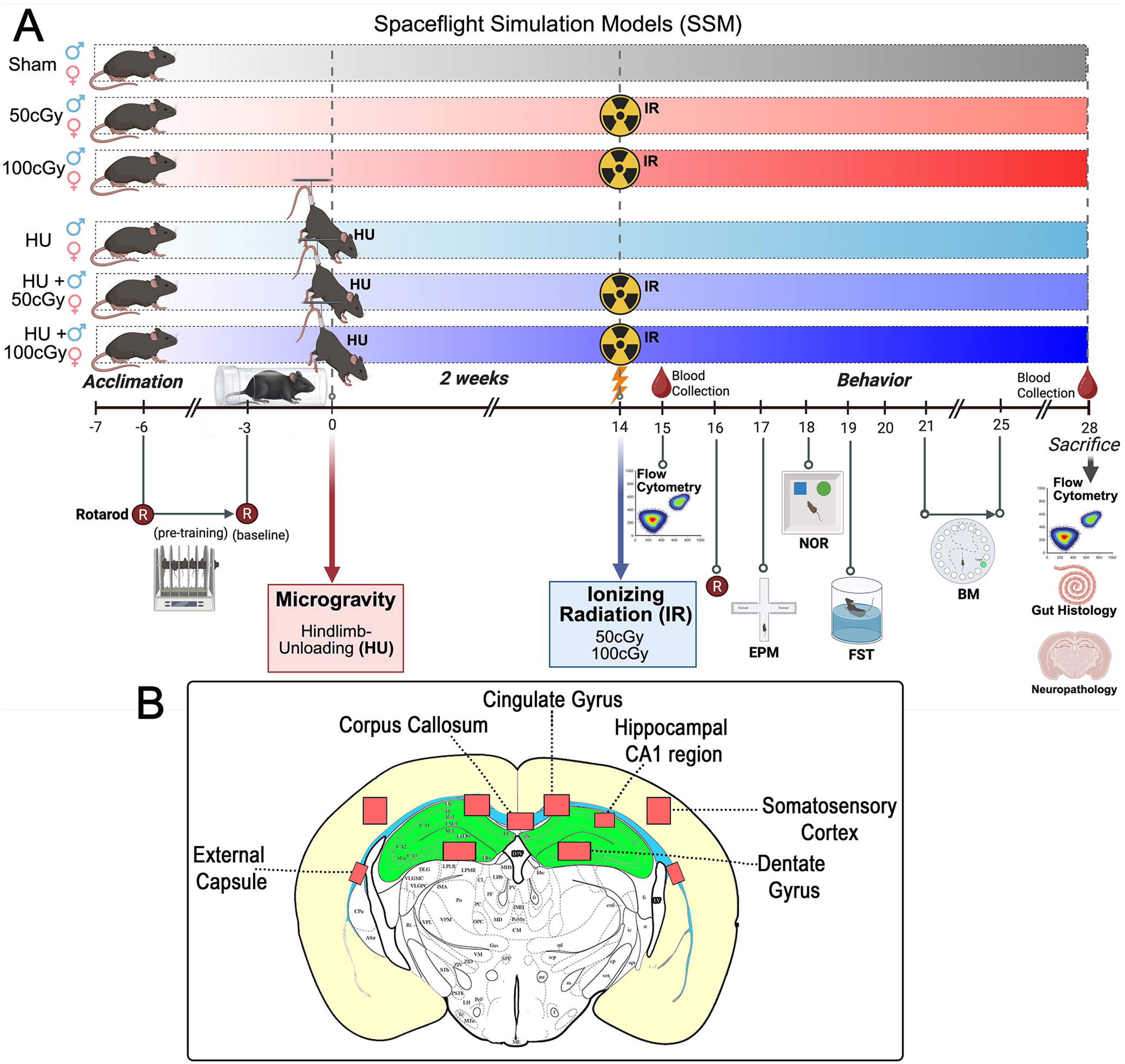
Experimental design for spaceflight simulation model exposure. **(A)** Adult male and female mice were assigned to six spaceflight simulation model (SSM) groups: Sham, ionizing radiation (IR; 50 or 100cGy), hindlimb unloading (HU; simulated microgravity), or combined HU+IR (HU+50cGy or HU+100cGy). Following acclimation, HU was initiated on day 0 and maintained for 14 days. On day 14, mice received a single dose of IR, as indicated. Behavioral testing was performed after exposure and included novel object recognition (NOR), the elevated plus maze (EPM), the forced swim test (FST), and the Barnes maze (BM). Rotarod performance was assessed during pre-training and baseline before SSM exposure and again after exposure. Blood was collected on day 15 and at sacrifice (day 28). Peripheral immune profiling was performed by flow cytometry at the interim time point and at the endpoint. At study completion, tissues were collected for flow cytometry, gut histology, and neuropathological analyses. **(B)** Schematic coronal brain section showing regions analyzed for neuropathology, including the corpus callosum, external capsule, cingulate gyrus, hippocampal CA1 region, somatosensory cortex, and dentate gyrus.

### 2.3. Submandibular blood collection

At 24 h post-irradiation, mice were anesthetized with isoflurane (2.5% induction) to collect blood for the flow cytometry studies. Submandibular blood was collected using a 5 mm animal lancet (Braintree Scientific, GR5MM). Approximately 200 µL of whole blood was collected into tubes containing 100 µL EDTA and maintained on ice. Samples were centrifuged at 400g for 15 min, and plasma was collected and aliquoted. The remaining cellular fraction was incubated in ACK lysis buffer for 5 min at room temperature, washed with PBS, and resuspended in freezing medium (90% FBS, 10% DMSO). Samples were stored at -80°C until further analysis.

### 2.4. Gut histology and intestinal immunostaining

#### 2.4.1. Tissue collection, fixation, processing, and sectioning

Segments of the small intestine, the jejunum (JE) and ileum (IL)and the large intestine, colon, were collected and fixed in 4% paraformaldehyde (PFA) for 24 h at 4°C. Tissues were then transferred to 70% ethanol and processed for paraffin embedding using a Shandon Excelsior ES tissue processor and embedded on a Shandon HistoCenter embedding system (Thermo Fisher Scientific) according to the manufacturer’s standard protocols. Paraffin blocks were sectioned at 5 µm and mounted onto glass slides.

#### 2.4.2. Hematoxylin and eosin (H&E) staining

Paraffin sections were deparaffinized in xylene and rehydrated through graded ethanols to water. Slides were stained with hematoxylin for 6 h at 60-70°C, rinsed in tap water, differentiated in 0.3% acid alcohol, rinsed again in tap water, and counterstained with eosin for 2 min. Sections were then rinsed, cleared in xylene, and coverslipped using a xylene-based mounting medium (Permount) and allowed to dry overnight.

#### 2.4.3. Alcian Blue staining

To assess goblet cell-associated mucin, sections were deparaffinized in xylene and rehydrated through graded ethanols to water. Slides were rinsed in distilled water and incubated with Alcian Blue solution for 30 min. Excess stain was removed by rinsing in tap water, followed by counterstaining with Nuclear Fast Red solution for 5 min. Sections were then rinsed, dehydrated through graded ethanols, cleared in xylene, and coverslipped as described above.

#### 2.4.4. Intestinal immunofluorescence staining

Paraffin sections were deparaffinized in xylene and rehydrated through graded ethanols to water. Antigen retrieval was performed using 10x Universal HIER antigen retrieval reagent (Abcam, ab208572) with heat-mediated retrieval (boiling) for 20 min. Sections were cooled in water, washed in 0.5% PBS-Triton X-100 (PBST), and blocked with normal goat serum for 1-2 h at room temperature. Slides were incubated with primary antibodies against ZO-1 (1:200; Invitrogen, 61-7300) and CD45 (1:250; BD Biosciences, 557390). After washing in PBST, sections were incubated with secondary antibodies anti-rabbit Alexa Fluor 488 (Thermo Fisher Scientific, A11034) and anti-rat Alexa Fluor 568 (Thermo Fisher Scientific, A11077) for 1 h at room temperature. Slides were washed in PBS and counterstained with DAPI (Thermo Scientific, PI62247) for 5 min. Sections were rinsed in deionized water and coverslipped.

### 2.5. Immune cell stimulation and flow cytometry

Cryopreserved blood leukocytes were retrieved from -80°C and rapidly thawed in a 37°C water bath. Cells were transferred to complete RPMI (cRPMI; RPMI supplemented with 10% FBS), and viable cells were quantified by Trypan blue exclusion using a hemocytometer. For each animal, 1-2 x 10^5^ viable cells were allocated per condition and split into unstimulated and stimulated fractions. Where applicable, additional aliquots were prepared for unstained, single-stain, and viability controls. Cells were resuspended in stimulation medium (RPMI containing 10% heat-inactivated FBS, 1% penicillin/streptomycin, 1% sodium pyruvate, 1% HEPES, 1% non-essential amino acids, and 1% 2-mercaptoethanol). Stimulated samples were incubated with a stimulation cocktail (BioLegend, 423304) for 5 h at 37°C. Following stimulation, cells were stained with a fixable viability dye (Thermo Fisher Scientific, 65-0863-18) and then surface-stained with antibodies against CD45 (Invitrogen, 368-0451-82), CD11b (Thermo Fisher Scientific, 36-601-1282), CD3 (Thermo Fisher Scientific, 367-0032-82), CD4 (BV650; BioLegend, 100469), CD8 (BUV661; BD Biosciences, 569186), CD19 (SparkUV387; BioLegend, 115586), CD25 (BV785; BioLegend, 102051), and NK1.1 (BUV495;

Thermo Fisher Scientific, 364-5941-82). Cells were washed and then fixed/permeabilized using the eBioscience Foxp3/Transcription Factor Staining Buffer Set (Thermo Fisher Scientific, 00-5523-00) with an overnight incubation at 4°C. Cells were subsequently stained intracellularly for FoxP3 (AF700; BioLegend, 126422), TNF-α (FITC; BioLegend, 506304), IL-4 (BV711; BioLegend, 504133), IFN-γ (BV605; BioLegend, 505840), and IL-10 (PE; Thermo Fisher Scientific, 12-7101-41). Data were acquired on a BD FACSymphony A5 SE cytometer using BD FACSDiva software. Unstained and single-stained controls were used for compensation and gating (**Supplementary Figure 1**). Flow cytometry standard (.fcs) files were exported and analyzed using FlowJo software.

### 2.6. Brain immunofluorescence

Following perfusion dissected brains were immersion-fixed in 4% PFA overnight at 4°C and cryoprotected in 30% sucrose in PBS for 24 h at 4°C. Coronal sections (15 µm) were cut on a cryostat at -20°C and either mounted onto glass slides and stored at -80°C or stored free-floating at -20°C in cryoprotectant solution (sucrose, sodium phosphate monobasic, sodium phosphate dibasic, ethylene glycol, and polyvinylpyrrolidone in distilled water). Mounted or free-floating sections were brought to room temperature and washed in PBS, followed by PBST. Sections were blocked in 10% normal goat serum for 1–2 h at room temperature and incubated with primary antibodies overnight at 4°C. Primary antibodies included MAP-2 (1:500; Cell Signaling Technology, 4542S), SMI-312 (1:1000; BioLegend, 837904), Iba-1 (1:500; Wako, 019-19741), GFAP (1:500; Millipore, MAB360), Lycopersicon esculentum (tomato) lectin 594 (1:1000; Vector Laboratories, DL11741), and claudin-5 (1:500; Abcam, ab15106). After PBST washes, sections were incubated with fluorophore-conjugated secondary antibodies for 1 h at room temperature in the dark (goat anti-mouse Alexa Fluor 488 [IgG], Thermo Fisher Scientific, A21121; goat anti-rabbit Alexa Fluor 647, A21244; goat anti-rabbit Alexa Fluor 488, A11034; goat anti-mouse Alexa Fluor 568, A11031). Sections were washed in PBS, counterstained with DAPI for 5 min, rinsed in deionized water, and coverslipped (mounted sections) or mounted onto slides prior to coverslipping (free-floating sections). Representative images were captured using the Leica confocal microscope at 40x magnification and a 6 μm z-stack.

### 2.7. Western Blot

Cortex brain tissue from each mouse was collected and gently homogenized on ice in a lysis buffer containing T-PER™ Tissue Protein Extraction Reagent (Thermo Fisher Scientific, 78510) and Protease Inhibitor Cocktail (Sigma-Aldrich, P8340). The mixtures were centrifuged at 13500 rpm for 20 min at 4°C, and the total protein in each sample was collected from the supernatants. The protein quantity of each sample was determined using the Pierce Bicinchoninic Acid (BCA) Protein Assay Kit (Thermo Fisher Scientific, 23227). Next, the samples were combined with 4x Laemmli sample buffer (Bio-Rad, 1610747) to achieve a final protein concentration of 60 μg, then heated at 100°C for 5 min. The samples were subsequently subjected to SDS-PAGE with a 4%-15% gradient for protein separation, followed by electro-transfer to polyvinylidene difluoride (PVDF) membranes (Bio-Rad, 1620177). The membranes were incubated with a blocking solution consisting of 5% w/v skim milk powder in PBS-Tween 20 buffer for 1 h at room temperature. Expression levels of the proteins to be analyzed were determined using antibodies against BDNF (1:1000, Abcam, ab108319), HSP90 (1:1000, Cell Signaling Technology, 4877), and HSP25 (1:1000, Stressgen, Spa-801), along with the corresponding HRP-conjugated secondary antibody. β-Actin antibody (Cell Signaling Technology, 3700) was used as a housekeeping protein. Immunoblots were visualized using Clarity Western ECL (Bio-Rad, 1705061) on a ChemiDoc MP imaging system (Bio-Rad) and subsequently quantified using a densitometer with ImageJ.

### 2.8. Experimental timeline and behavioral testing

A schematic of the Spaceflight Simulation Model (SSM) experimental design and timepoints is shown in **Fig. 1**. Following acclimation (7 days; day -7 to 0), mice were assigned to sham, ionizing radiation (IR; 50 or 100 cGy), HU, or combined HU+IR groups. HU (or sham HU) was initiated on day 0 and maintained for 14 days. On day 14, mice received whole-body X-ray irradiation (0, 50, or 100 cGy) and were reloaded immediately thereafter. Submandibular blood samples for immune analyses were collected 24 h post-irradiation (day 15) and used for flow cytometry. Beginning on day 16, mice underwent behavioral testing, and terminal blood and tissue collection was performed at sacrifice on day 28. All behavioral testing was conducted by an experimenter blinded to group assignment.

#### 2.8.1. Rotarod

Sensorimotor function was assessed using an accelerating rotarod (Ugo Basile, Gemonio, Italy). Mice underwent rotarod pre-training on day -6 and baseline assessment on day -3. Post-simulation performance was assessed beginning on day 16. For each session, mice were placed on the stationary rod for 30 s, after which the rod accelerated from 4 to 40 rpm over 5 min. Trials ended when the mouse fell or when the 5-min cutoff was reached. Latency to fall was recorded and averaged per mouse.

#### 2.8.2. Elevated plus maze

Anxiety-like behavior was evaluated using the elevated plus maze (EPM) at day 3 post-SSM. The apparatus was elevated 75 cm above the floor and consisted of two open arms (50x10 cm) and two closed arms (50x10x40 cm) extending from a central platform. Mice were placed on the central platform and allowed to explore for 10 min. Time spent in open and closed arms was quantified.

#### 2.8.3. Forced swim test

Depressive-/stress-coping-like behavior was assessed using the forced swim test (FST) at day 5 post-SSM. Mice were placed in a transparent cylindrical tank (30 cm height x 20 cm diameter) filled with room-temperature water to a depth of 15 cm and recorded for 6 min. Immobility (lack of active escape-directed movements) was quantified from video recordings. After testing, mice were dried and placed in a warmed recovery cage lined with paper towels on a heating pad.

#### 2.8.4. Novel object recognition

Recognition memory was assessed using the novel object recognition (NOR) task at day 4 post-SSM. Mice were habituated to an empty arena (22x44 cm) for 5 min. During the acquisition phase, mice explored two identical objects for 5 min. In the test phase (5 min), one familiar object was replaced with a novel object. Time spent exploring each object was recorded and used to derive recognition performance (e.g., discrimination index).

#### 2.8.5. Barnes maze

Spatial learning and memory were assessed using the Barnes maze (BM) at day 7-11 post-SSM. The circular platform (122 cm diameter) contained 40 equidistant holes (5 cm diameter), with one target hole leading to an escape box. Distal visual cues (4) were positioned around the maze, and white noise (90 dB) was used to mask background noises. Trials were initiated by placing the mouse in an opaque start chamber at the center of the maze; lighting was adjusted to ∼1000 lux as an aversive stimulus at trial start. Mice were allowed up to 3 min to locate the escape box. Training consisted of two trials per day for four consecutive days, followed by a probe trial on day 5 in which the escape box was removed and the former target hole was blocked. Probe performance was quantified as time spent in the target zone relative to other zones, and search strategy was classified as spatial, spatial/sequential, sequential, or mixed/random.

### 2.9. Quantitative image analysis and statistics

Alcian Blue-stained intestinal sections were digitized in their entirety using an Olympus VS200 slide scanner. Morphometric measurements were performed in ImageJ (v2.14.0/1.54f) for JE and IL regions. Alcian Blue–positive area (% area) was quantified in QuPath (v0.6.0, x64). Immunofluorescence-stained gut and brain sections were imaged using the VS200 slide scanner, and quantitative analyses were performed in QuPath (v0.6.0, x64). For brain region-of-interest (ROI) analyses, ROIs were annotated in QuPath using neuroanatomical landmarks and included the somatosensory cortex (SSC), cingulate gyrus (CG), corpus callosum (CC), external capsule (EC), hippocampal CA1, dentate gyrus (DG), and hypothalamic nuclei (paraventricular, dorsomedial, and arcuate). To account for regional heterogeneity in vascular density and staining intensity across sections, a vessel-centric, pixel-based analysis pipeline was implemented in QuPath (**Supp. Fig. 2A**). Blood vessels were detected using a supervised pixel classifier trained on lectin-mCherry signal. Perivascular compartments were defined by expanding vessel objects by 10µm and subtracting the vessel core, generating a perivascular “shell” for each vessel. IgG signal was quantified within perivascular shells and normalized to vessel area within each ROI. IgG-positivity thresholds were determined per image using background-intensity statistics to reduce bias from intersession variability. Statistical analyses were performed in GraphPad Prism (v10.6.1). Two-way ANOVA was used for factors of interest (e.g., HU and IR, with sex included where indicated), followed by Benjamini-Hochberg correction for multiple comparisons and Tukey’s multiple comparisons test. Adjusted p-values ≤ 0.05 were considered statistically significant. Data are presented with 95% confidence intervals (95% CI) around the mean based on the standard error of the mean (SEM).

**Figure 2.**
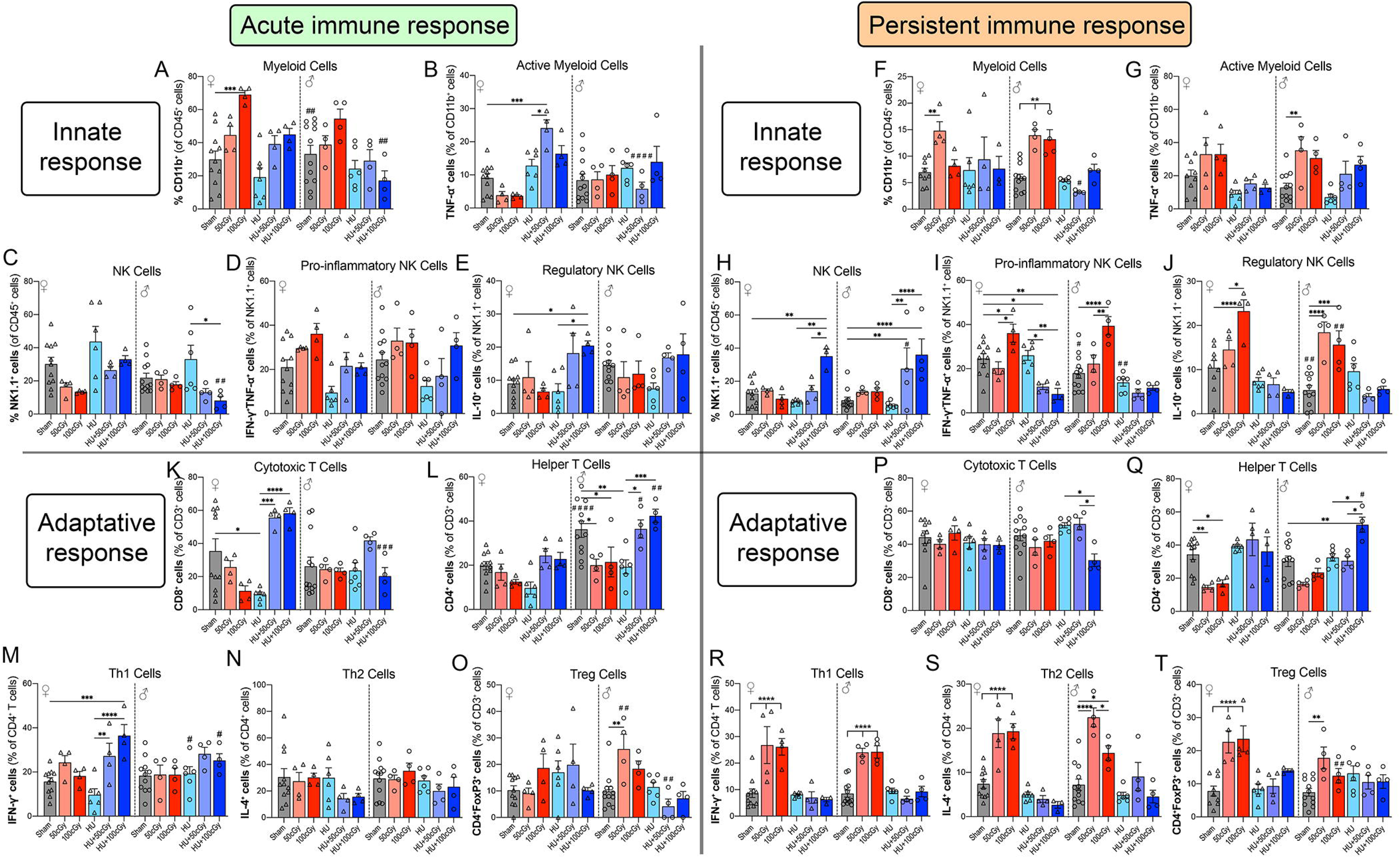
Acute and persistent peripheral immune responses after spaceflight simulation model exposure. Flow cytometric analysis of peripheral immune cell populations was performed at acute and persistent time points following exposure to hindlimb unloading (HU), ionizing radiation (IR; 50 or 100cGy), or combined HU+IR. For the innate immune response, the proportions of myeloid cells (A, F), active myeloid cells (B, G), NK cells (C, H), pro-inflammatory NK cells (D, I), and regulatory NK cells (E, J) were quantified. For the adaptive immune response, the proportions of cytotoxic T cells (K, P), helper T cells (L, Q), Th1 cells (M, R), Th2 cells (N, S), and regulatory T cells (Treg) (O, T) were quantified. Panels A–E and K–O show the acute immune response, whereas panels F-J and P-T show the persistent immune response. Cell populations are presented as percentages of their corresponding parent populations, as defined by the flow cytometry gating strategy. Adaptive and innate immune subsets were identified using lineage-specific surface and intracellular markers, with representative gating shown in the Supplementary Figure 1. Data are shown as mean ± SEM. *p<0.05, **p < 0.01, ***p<0.001, ****p<0.0001. Statistical significance in sex differences is shown as #p<0.05, ##p<0.01, ####p<0.0001.

## 3. RESULTS

### 3.1. Simulated spaceflight induces acute and persistent immune alterations in innate and adaptive compartments

To determine whether simulated spaceflight conditions alter systemic immune responses, peripheral immune cell populations were analyzed by flow cytometry following PMA/ionomycin stimulation of isolated leukocytes at two time points: *<u>an acute phase</u>* and *<u>a persistent phase</u>* after exposure. Immune profiling included both innate immune populations (myeloid and NK cells) and adaptive immune populations (T cell subsets) (**Fig. 2 and Supp. Fig. 1**).

#### Innate immune responses

Innate immune populations exhibited pronounced alterations following simulated spaceflight exposures. During the acute phase, myeloid cells (CD11b⁺) increased significantly in female mice exposed to 100cGy radiation (***p<0.001), indicating rapid activation of innate immune pathways (**Fig. 2A**). Activated myeloid cells (TNFα⁺CD11b⁺) were particularly elevated in female groups exposed to combined HU and radiation compared to sham (***p<0.001), HU-only (*p<0.05), and males of the same exposure (###p<0.001) (**Fig. 2B**), suggesting sex-specific enhanced inflammatory signaling under combined stress conditions. Natural killer (NK) cell populations were also affected. Total NK cells (NK1.1⁺) have increased trends following HU exposure (both sexes) and combined exposures (females) (**Fig. 2C**), while pro-inflammatory NK cells (IFNγ⁺/TNFα⁺) exhibited exposure-dependent changes (**Fig. 2D**). Regulatory NK cells (IL-10⁺/NK1.1⁺) were elevated in several combined exposure groups (*p<0.05) (**Fig. 2E**), suggesting alterations in NK cell functional balance. At the *persistent time point*, innate immune alterations were still evident. Myeloid cells remained elevated in radiation exposure groups (**p<0.01) (**Fig. 2F**), while activated myeloid populations continued to display increased inflammatory signaling (**p<0.01 50cGy males) (**Fig. 2G**). NK cell populations and their functional subsets showed sustained dysregulation, with increases in total NK cells in females following HU+100cGy (**p<0.01) and males post HU+100cGy (****p<0.0001) and HU+50cGy (**p<0.01) compared to sham and HU+50cGy females (#p<0.05). Pro-inflammatory and regulatory NK cell phenotypes are increased in both sexes following radiation exposures (*p<0.05, **p<0.01, ***p<0.001, ****p<0.0001). Interestingly, NK population subsets are decreased following HU+IR exposures, showing significance in female pro-inflammatory NK subsets (*p<0.05, **p<0.01) (**Fig. 2I-J**).

#### Adaptive immune responses

Analysis of adaptive immune populations revealed significant changes across cytotoxic T cells (Tc, CD8⁺), helper T cells (Th, CD4⁺), and their functional subsets. During the *acute phase*, cytotoxic T cell frequencies were altered in female groups, with the most prominent shifts observed following HU (decreased *p<0.05) compared to sham, and significant increases in HU+IR groups compared to HU-only (***p<0.001, ****p<0.0001) (**Fig. 2K**). Helper T cells also exhibited exposure-dependent alterations, with decreases detected in IR and HU independent exposure male groups (*p<0.05, **p<0.01), indicating early adaptive immune loss. Male Th responses following HU+IR are also significantly elevated compared to HU (*p<0.05, ***p<0.001) and compared to females after HU+50cGy (#p<0.05) and HU+100cGy (##p<0.01) (**Fig. 2L**). Functional polarization of helper T cells was also affected. Th1 cells (IFNγ⁺CD4⁺) showed increases following combined exposures in females compared to sham (***p<0.001), HU (****p<0.0001), and same-exposure males (#p<0.05) (**Fig. 2M**). Th2 cells (IL-4⁺ CD4⁺) exhibited minimal responses across groups (**Fig. 2N**). In contrast, regulatory T cells (Treg; FOXP3⁺CD4⁺) were significantly elevated (**p<0.01) in males after 50cGy compared to sham and 50cGy females (##p<0.01). Sex-specific shifts in trends were also present after simulated spaceflight stressors (**Fig. 2O**), suggesting an early compensatory immunoregulatory response. At the *persistent time point*, adaptive immune alterations remained evident. Cytotoxic T cell populations were unchanged compared to sham (**Fig. 2P**), but helper T cells decreased in females following IR (*p<0.05, **p<0.01) and increased in males after HU+100cGy conditions (**p<0.01) compared to sham and HU+100cGy exposed females (#p<0.05) (**Fig. 2Q**). Th1 and Th2 populations exhibit robust elevations after radiation exposure in both sexes (****p<0.0001), despite decreases in parent Th populations (**Fig. 2R-S**). Regulatory T cells also remained elevated in males post 50cGy (**p<0.01) and significantly increase in females following IR (****p<0.0001) (**Fig. 2T**), suggesting prolonged immune modulation following simulated spaceflight. Together, these findings demonstrate that simulated spaceflight induces sexually dimorphic acute and persistent immune alterations, characterized by innate immune activation and sustained modulation of adaptive immune populations. These immune changes suggest systemic inflammatory and immunoregulatory responses that may contribute to downstream neurological and physiological alterations observed after simulated spaceflight exposure.

### 3.2. Spaceflight simulation alters axonal structure across cortical and white-matter regions

Axonal and dendritic integrity are essential for maintaining neuronal connectivity, signal transmission, and protection against neurodegenerative processes^47^. To evaluate structural neuronal alterations induced by simulated spaceflight conditions, we assessed MAP-2, a dendritic marker, and SMI-312, a pan-axonal neurofilament marker associated with axonal structure and injury. Immunofluorescence analyses were performed in the somatosensory cortex (SSC), while axonal integrity was additionally quantified in major white-matter tracts including the corpus callosum (CC), cingulate gyrus (CG), and external capsule (EC) (**Fig. 3**). In the SSC, MAP-2 staining revealed significant reductions in expression following several simulated spaceflight exposures compared with sham controls (**Fig. 3F**). Radiation had a dose-dependent impact (*p<0.05 50cGy, *p<0.05 or **p<0.01 100cGy) decrease in MAP-2 signal, yet the addition of HU dampens MAP-2 signal loss. Combined exposures partially attenuated the radiation-induced loss of MAP-2. Analysis of SMI-312 immunoreactivity demonstrated alterations in axonal structure across exposure groups (**Fig. 3G-H**). In the somatosensory cortex, the female 50cGy and HU+50cGy group showed a significant increase in SMI-312 signal (*p<0.05), suggesting axonal remodeling or injury, whereas the other exposure groups showed similar trends that did not reach statistical significance. To determine whether axonal alterations extended to white-matter regions, SMI-312 expression was quantified in the CC, CG, and EC (**Fig. 3I-K**). Radiation exposure produced greater and more consistent changes in axonal neurofilament signal compared to sham (**p<0.01 50cGy, ***p<0.001 100cGy) than HU alone, with evidence of dose-dependent effects across regions. Combined HU and radiation exposure also altered SMI-312 expression in the CG (**p<0.01 males) and EC (*p<0.05 females), indicating that simulated microgravity and radiation together can influence axonal structure within major interhemispheric and cortical projection pathways **(Fig. 3J-K**). Furthermore, we measured brain-derived neurotrophic factor (BDNF) levels in the brain tissue by western blot (**Fig. 3L-M**). HU+100cGy exposure significantly elevated BDNF levels in male mice compared to sham (***p<0.001), indicating a compensatory neuroprotective response, while no significant changes were observed in female mice after several simulated spaceflight exposures. Notably, BDNF levels in HU+100cGy-exposed males were significantly higher than those observed in HU+100cGy-exposed females (#p<0.05). Together, these findings indicate that simulated spaceflight conditions disrupt neuronal structural integrity, with radiation producing the most pronounced effects and combined exposures contributing to region-specific alterations in axonal and dendritic architecture. These structural changes suggest potential impairment of neuronal connectivity and information transfer within cortical and white-matter circuits following simulated spaceflight exposure.

**Figure 3.**
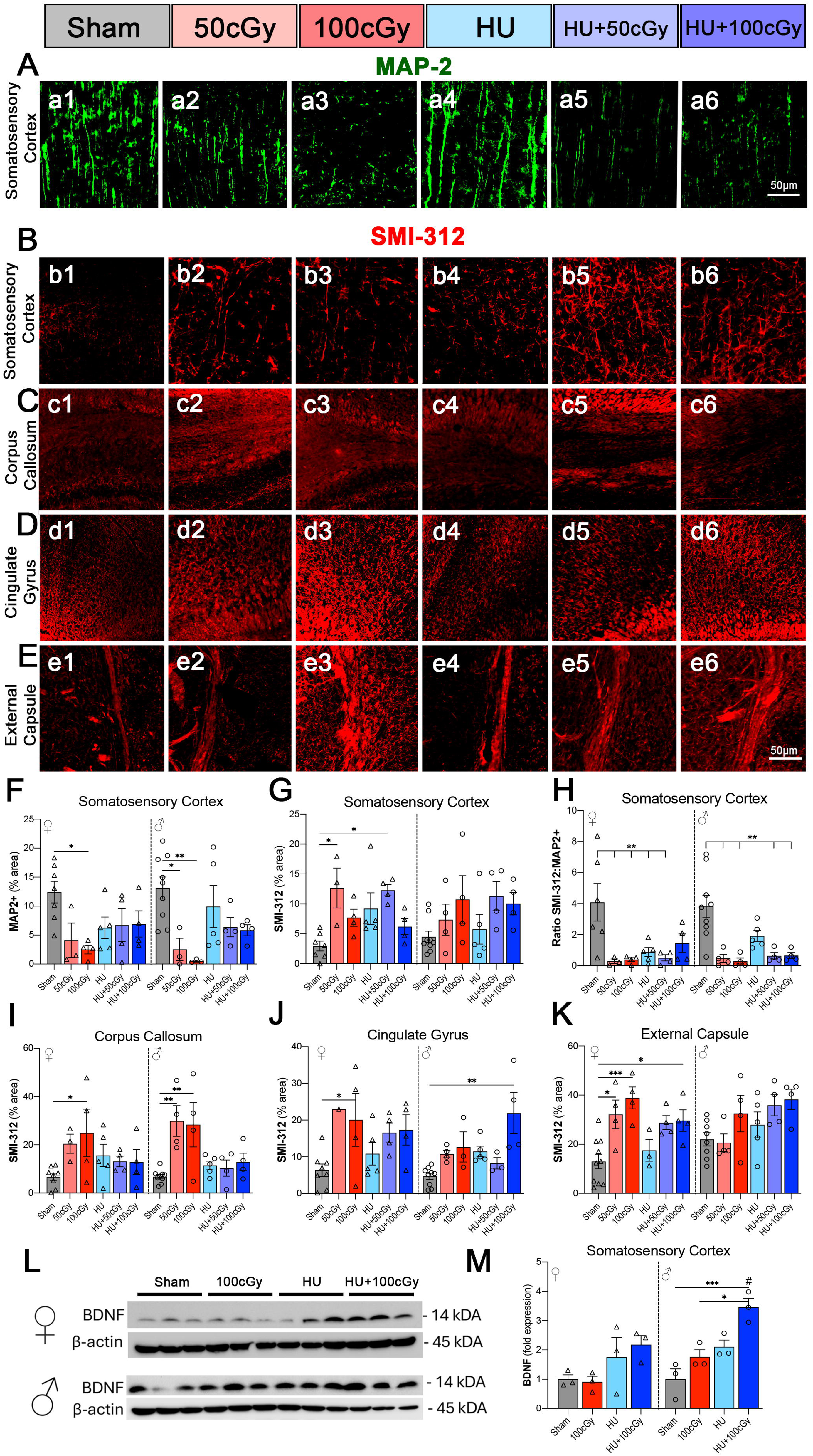
Axonal integrity and neurotrophic signaling following spaceflight simulation model exposure. Representative immunofluorescence images and quantitative analyses of axonal and neuronal markers in selected brain regions following exposure to hindlimb unloading (HU), ionizing radiation (IR; 50 or 100cGy), or combined HU+IR. (A-E) Representative images showing MAP-2 (green; dendritic marker) and SMI-312 (red; pan-axonal neurofilament marker) immunoreactivity in the somatosensory cortex (A-B), corpus callosum (C), cingulate gyrus (D), and external capsule (E) across experimental groups (Sham, 50cGy, 100cGy, HU, HU+50cGy, HU+100cGy). Scale bars: 50μm. (F-H) Quantification of MAP-2⁺ dendritic area (F), SMI-312⁺ axonal signal intensity (G), and relative SMI-312 density (H) in the somatosensory cortex. (I-K) Quantification of SMI-312⁺ axonal signal intensity in white matter regions, including the corpus callosum (I), cingulate gyrus (J), and external capsule (K). (L) Representative western blot analysis of brain-derived neurotrophic factor (BDNF) protein expression in brain tissue from female and male mice following exposure to Sham, IR (100cGy), HU, or HU+100cGy conditions. β-actin was used as a loading control. (M) Quantification of BDNF protein levels normalized to β-actin and expressed relative to Sham controls, stratified by sex. Representative images depict the largest response among males and females. Data are presented as mean ± SEM. **p<0.01, ***p<0.001. Statistical significance in sex differences is shown as #p<0.05. Statistical significance in sex differences is shown as #p<0.05.

### 3.3. Simulated spaceflight promotes microglial activation and astrogliosis across hippocampal and white-matter regions

To determine whether simulated spaceflight conditions induce sustained neuroinflammation, we assessed microglial activation and astrocyte reactivity by immunofluorescence staining for Iba-1 and GFAP across four brain regions of interest: CA1, dentate gyrus (DG), CG, and CC (**Fig. 4A-D**). Quantitative analyses were performed separately for female and male mice. Within the hippocampus, both sexes activated significant increases in microglial expression and astrogliosis following combined HU+IR exposure compared with sham controls (**Fig. 4E-H**). In the CA1 region, Iba-1 expression increased in a dose-dependent manner, with significant elevations observed after HU+50cGy exposure (females **p<0.01, males *p<0.05) and the largest increases detected in the HU+100cGy group (****p<0.0001) (**Fig. 4E**). GFAP immunoreactivity in CA1 also increased significantly in combined exposure groups, indicating enhanced astrocyte activation (*p<0.05, **p<0.01) (**Fig. 4F**). Similar trends were observed in the DG, where both Iba-1 and GFAP expression were significantly elevated following HU+100cGy exposure (*p<0.05, **p<0.01), indicating persistent inflammatory responses in hippocampal circuits (**Fig. 4G-H**). Region-specific effects were also observed in cortical and white-matter structures. In the CG, astrocyte activation was not significantly altered across exposure groups (**Fig. 4J**). However, microglial expression increased in females following independent HU (*p<0.05) and 100cGy radiation (**p<0.01) exposures compared with sham, whereas males showed significant increases only after the combined HU+100cGy exposure (*p<0.05) (**Fig. 4I**). These findings suggest sex-dependent regional sensitivity to simulated microgravity and radiation. The CC, a major white-matter tract, displayed a similar pattern of inflammatory activation. Both female and male mice exhibited significant increases in Iba-1 expression following HU+100cGy exposure (*p<0.05), indicating microglial activation in white-matter regions (**Fig. 4K**). In contrast, GFAP expression increased significantly only in males following HU+100cGy exposure (**p<0.01), suggesting sex-specific astrocytic responses within this tract (**Fig. 4L**). To further evaluate molecular stress responses associated with neuroinflammation, we measured heat shock proteins HSP90 and HSP25 by western blot (**Fig. 4M-O**). HSP90 expression was reduced after HU (*p<0.05) and combined HU+100cGy (**p<0.01) exposures in females but increased following 100cGy (*p<0.05) conditions in males. Female HSP25 levels were significantly elevated following HU+100cGy exposure compared to HU (**p<0.01), whereas HU was significantly reduced compared to sham (**p<0.01), consistent with activation of cellular stress-response pathways. Together, these findings demonstrate that simulated spaceflight induces region-specific neuroinflammation characterized by microglial activation and astrocytic reactivity, particularly within hippocampal and white-matter regions. The most pronounced effects occurred under combined microgravity and radiation conditions, with notable sex-dependent differences, suggesting that females may exhibit heightened inflammatory sensitivity to simulated microgravity stressors.

**Figure 4.**
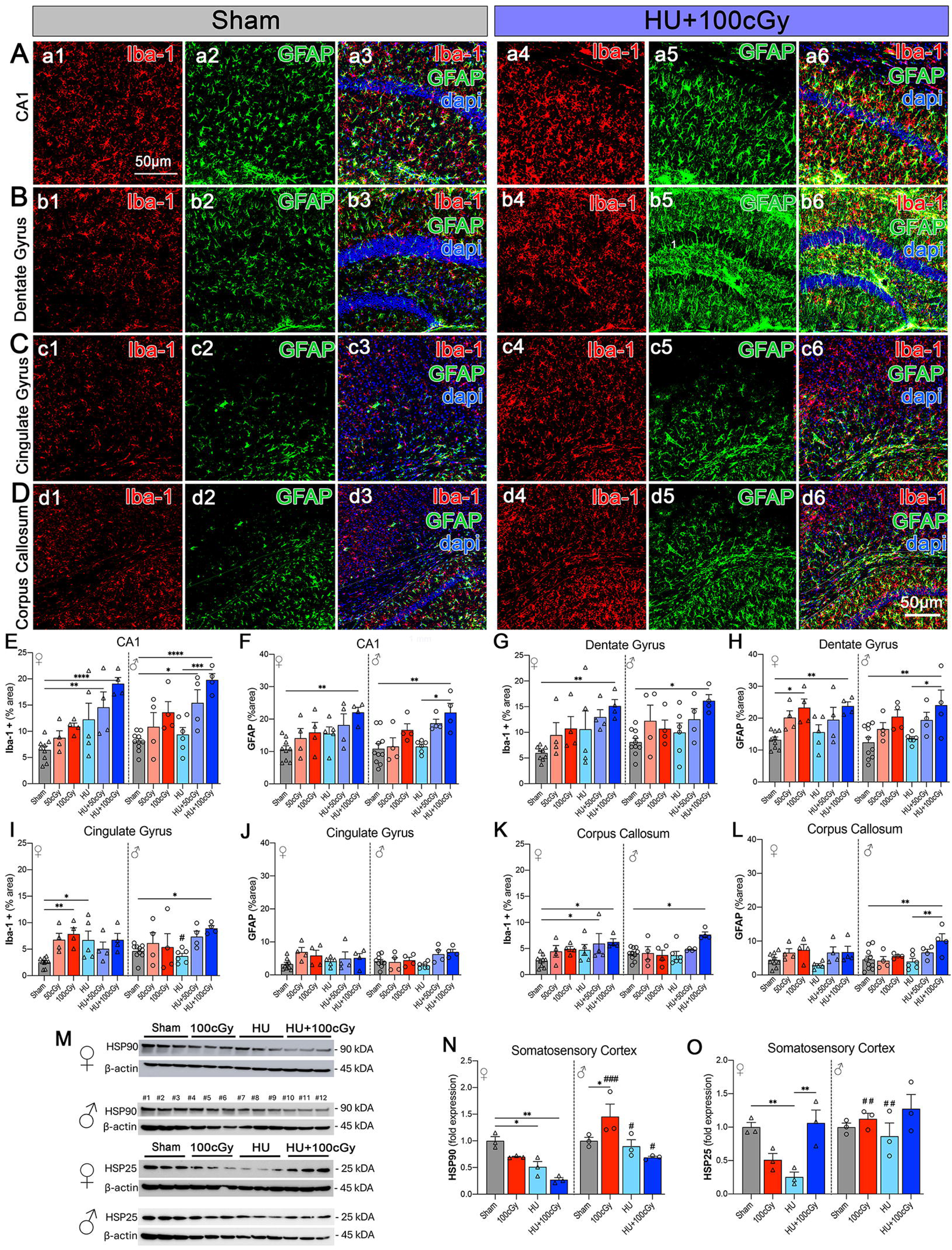
Microglial activation and astrogliosis following spaceflight simulation model exposure. Representative immunofluorescence images and quantitative analyses of microglial and astrocytic markers in selected brain regions after exposure to hindlimb unloading (HU), ionizing radiation (IR; 50 or 100cGy), or combined HU+IR. (A-D) Representative images of Iba-1 (red; microglia), GFAP (green; astrocytes), and DAPI (blue; nuclei) in the CA1 hippocampal region (A), dentate gyrus (B), cingulate gyrus (C), and corpus callosum (D) for Sham and HU+100cGy groups. Merged images show colocalized staining within each region of interest. Scale bars: 50μm. (E-L) Quantification of Iba-1⁺ microglial area (%) and GFAP⁺ astrocytic area (%) across experimental groups in the CA1 region (E-F), dentate gyrus (G-H), cingulate gyrus (I-J), and corpus callosum (K-L). Data are shown separately for female and male mice. (M) Representative western blot analysis of HSP90 and HSP25 protein expression in brain tissue from female and male mice following Sham, IR (100cGy), HU, and HU+100cGy exposure. β-actin served as the loading control. (N-O) Quantification of HSP90 (N) and HSP25 (O) protein levels normalized to β-actin and expressed relative to Sham controls. Representative images depict the largest response among males and females. Data are presented as mean ± SEM, with each point representing an individual animal. Statistical analysis was performed using two-way ANOVA with appropriate multiple-comparisons testing, with statistical significance indicated: *p<0.05, **p<0.01, ***p<0.001, ****p<0.0001. Statistical significance in sex differences is shown as ##p<0.01, ####p<0.0001.

### 3.4. Simulated spaceflight disrupts blood–brain barrier integrity and promotes IgG leakage

Given that BBB dysfunction has been reported after spaceflight exposure^17,18^, we evaluated BBB structural integrity and permeability by immunofluorescence staining of vascular endothelium (lectin), the tight-junction protein claudin-5 (Cldn-5), and endogenous immunoglobulin G (IgG) as an indicator of vascular leakage^48^ (**Fig. 5** and **Supp. Fig. 2**). Analyses were performed in multiple brain regions including the DG, CA1 hippocampal region, CG, and SSC. Assessment of BBB structure revealed significant reductions in Cldn-5 signal within lectin-positive vessels, indicating impaired tight-junction integrity following simulated spaceflight exposures (**Fig. 5B, E, H, K**). In the DG, claudin-5 area was significantly reduced after 50cGy, 100cGy and HU+100cGy exposures (*p<0.05) in females, with males having significant reductions after HU and HU+100cGy (*p<0.05) (**Fig. 5B**). Similar reductions were observed in the CA1 region, where Cldn-5 expression decreased significantly across multiple exposure groups, particularly following radiation and combined exposures (**Fig. 5E**). In the CG and somatosensory cortex, tight-junction disruption was also evident, with significant decreases in claudin-5 expression following radiation (*p<0.05, **p<0.01, ****p<0.0001) exposure and combined HU+IR (**p<0.01, ****p<0.0001) conditions in both sexes (**Fig. 5H, K**). Across regions, radiation exposure produced the most consistent loss of tight-junction signal. HU alone produced more modest effects in females, yet induced significant reductions in males in each region compared to sham (*p<0.05 DG, and CG; **p<0.01 CA1 and SSC) and in the SSC compared to HU females (###p<0.001). To determine whether structural BBB disruption resulted in increased permeability, we quantified IgG leakage into the brain parenchyma (**Fig. 5C, F, I, L**). Endogenous IgG is normally confined to the vasculature and efficiently cleared by endothelial cells; thus, its presence within the parenchyma reflects BBB permeability^48^. IgG signal within the parenchymal space was measured within a defined radius (10μm) surrounding lectin-positive vessels and normalized to total vascular area (**Supp. Fig. 2A**). While endogenous IgG is detected at low levels in the brain vasculature, it is perceived to be digested by lysosomes and blocked by the low endocytic activity of brain endothelial cells^49^. Loss of BBB structure has been proven to allow IgG infiltration into the brain parenchyma^49^. Consistent with the observed disruption of tight junction integrity, IgG infiltration modestly increased across exposures, only significant in the SSC after HU+100cGy (*p<0.05 females, **p<0.01 males) (**Supp. Fig. 2F-I**). IgG leakage increased across several brain regions following simulated spaceflight exposures. In the DG, females exhibited significantly greater IgG leakage than males following HU (**p<0.01) and 50cGy (****p<0.0001) compared to sham and 50cGy males (###p<0.001) (**Fig. 5C, Supp. Fig 2D**). In the CA1 region, IgG leakage was elevated across multiple exposures in males, with significant increases observed particularly after 100cGy radiation (***p<0.001) and HU (***p<0.001) exposure (**Fig. 5F**). Both sexes showed increased leakage after radiation exposure, although the magnitude and statistical significance varied between sexes and exposure conditions, with significantly higher IgG levels in males compared to females following 100cGy (#p<0.05) (**Fig 5F, Supp. Fig. 2E**). Regional differences were also evident in cortical areas. In the CG, both female and male mice showed increased IgG leakage following 100cGy radiation (*p<0.05, **p<0.01), while HU exposure produced stronger leakage responses in females compared to sham (****p<0.0001) and HU males (##p<0.01) (**Fig. 5I, Supp. Fig. 2B**). In the SSC, IgG leakage was significantly elevated following HU exposure in females (****p<0.0001) and males (*p<0.05) compared to sham, although IgG leakage in females was still significantly higher than males (###p<0.001) (**Fig. 5L, Supp. Fig. 2C**). Across regions, HU exposure often produced strong IgG leakage despite smaller changes in Cldn-5 expression, suggesting that microgravity-related physiological changes may affect BBB permeability through mechanisms beyond tight-junction disruption alone. Together, these findings demonstrate that simulated spaceflight compromises BBB integrity and promotes vascular leakage across multiple brain regions. Radiation exposure produced substantial tight-junction disruption, whereas simulated microgravity was associated with pronounced IgG extravasation. Notably, female mice frequently exhibited greater BBB permeability than males, indicating sex-dependent vulnerability to spaceflight-associated vascular dysfunction.

**Figure 5.**
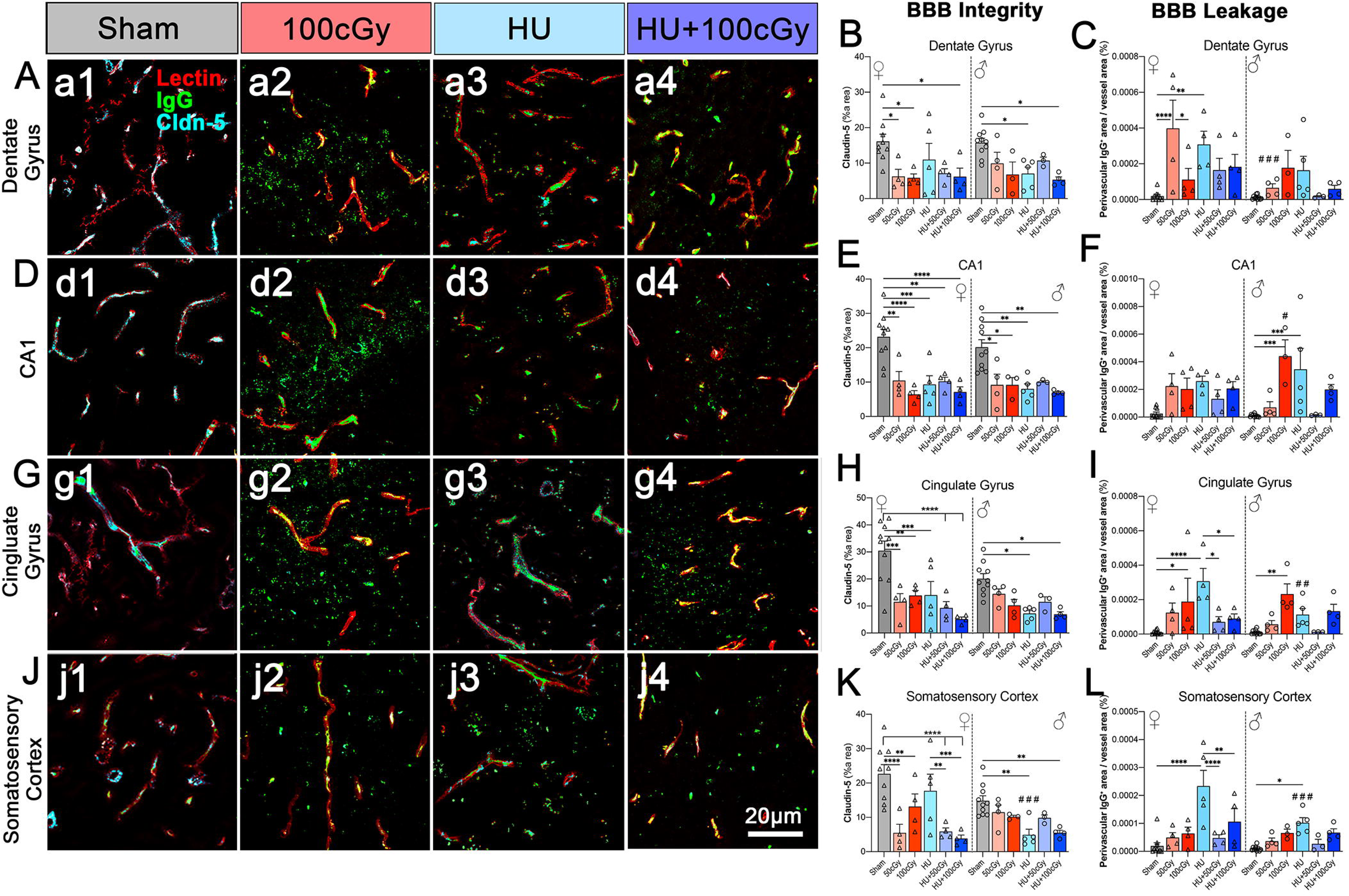
Blood-brain barrier integrity and leakage following exposure to a spaceflight simulation model. (A) Representative immunofluorescence images of blood–brain barrier (BBB) components in the dentate gyrus, CA1, cingulate gyrus, and somatosensory cortex from mice exposed to Sham, ionizing radiation (IR; 100cGy), hindlimb unloading (HU), or combined HU+100 cGy. Sections were stained for lectin (red; blood vessels), Claudin-5 (Cln-5) (blue; endothelial tight junction marker), and immunoglobulin G (IgG) (green, immune leakage marker). Merged images illustrate BBB structural organization across experimental groups. Scale bar: 20 μm. (B, D, H, K) Quantification of BBB integrity, measured as the relative area/intensity of tight junction markers within each region of interest in the dentate gyrus (B), CA1 (D), cingulate gyrus (H), and somatosensory cortex (K). (C, F, I, L) Quantification of BBB leakage, assessed by disruption and reduction of tight junction signal, in the dentate gyrus (C), CA1 (F), cingulate gyrus (I), and somatosensory cortex (L). Representative images depict the largest response among males and females. Data are shown separately for female and male mice. Values represent mean ± SEM, with each dot representing an individual animal. Statistical significance was determined using two-way ANOVA followed by multiple-comparisons testing, with statistical significance indicated: *p<0.05, **p<0.01, ***p<0.001, ****p<0.0001. Statistical significance in sex differences is shown as #p<0.05, ##p<0.01###p<0.001.

### 3.5. Simulated spaceflight disrupts small-intestinal architecture and alters mucin production in a sex-dependent manner

To evaluate whether simulated spaceflight affects intestinal morphology, Alcian blue staining was performed in the JE and IL to assess villus-crypt architecture and mucin production (**Fig. 6A-F**). Intestinal morphology dynamically responds to physiological stress and inflammation, where shortened villi and enlarged crypts represent hallmarks of intestinal injury^50,51^. The villus:crypt ratio provides a normalized indicator of intestinal inflammation, with ratios below 5:1 in the JE and 3:1 in the IL reflecting compromised mucosal architecture. Quantification of villus-crypt ratios revealed significant reductions in both the JE and IL across simulated spaceflight exposures compared with sham controls (**Fig. 6C-D**). In the JE, both female and male mice showed decreased ratios following radiation exposure, with reductions of approximately 1.2-1.5-fold relative to sham. The effect was more pronounced with HU alone, in which villus-crypt ratios decreased by 1.6-1.7-fold, indicating stronger microgravity-associated morphological disruption. Combined exposures (HU+IR) produced the largest decreases (****p<0.0001), reaching 1.6 - 2-fold reductions, suggesting additive effects of radiation and simulated microgravity on intestinal architecture. Similar patterns were observed in the IL, where villus-crypt ratios decreased significantly across exposure groups in both sexes, further supporting the presence of small-intestinal inflammation.

**Figure 6.**
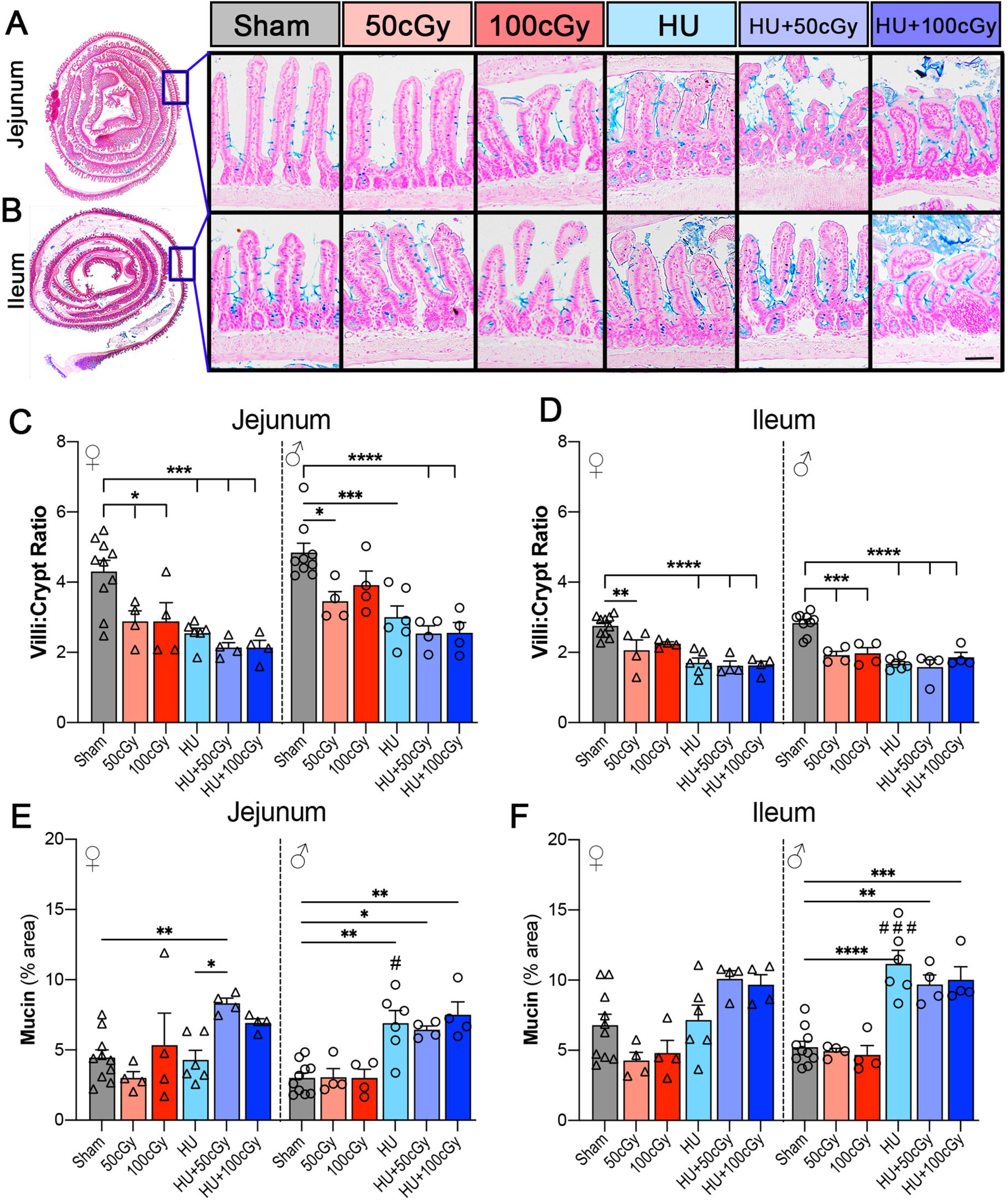
Spaceflight simulation alters small intestinal morphology and mucin content in a sex-dependent manner. (A-B) Representative histological sections of the jejunum (A) and ileum (B) from mice exposed to Sham, ionizing radiation (IR; 50 or 100cGy), hindlimb unloading (HU; simulated microgravity), or combined HU+IR (HU+50cGy and HU+100cGy). Low-magnification images (left) indicate the region analyzed, and higher-magnification images (right) show villus-crypt architecture and mucosal morphology across experimental groups. (C-D) Quantification of the villus:crypt ratio in the jejunum (C) and ileum (D). (E-F) Representative images depict the largest response among males and females. Quantification of mucin-positive area (% area) in the jejunum (E) and ileum (F). Data are shown separately for female and male mice. Bars represent mean ± SEM, and symbols indicate individual animals. Statistical significance was determined by two-way ANOVA with appropriate multiple-comparisons testing, with statistical significance indicated: *p<0.05, **p<0.01, ***p<0.001, ****p<0.0001. Statistical significance in sex differences is shown as #p<0.05, ###p<0.001. HU, hindlimb unloading; IR, ionizing radiation.

Because the intestinal mucin layer represents the first line of immune defense against luminal microbes and toxins^52^, we next quantified mucin production by measuring the percent area of Alcian blue-positive staining (**Fig. 6E-F**). In the JE, both sexes exhibited significant increases in mucin production after HU+50cGy exposure compared with sham (**p<0.01 in females; *p<0.05 in males). Male mice also displayed significantly elevated mucin production following HU alone and HU+100cGy exposure compared with sham (**p<0.01), with mucin levels also significantly higher than those observed in HU-exposed females (#p<0.05). In the IL, sex-dependent differences were more pronounced. Female mice showed reduced mucin production following radiation exposure, followed by significant increases in the JE after combined HU+IR exposures relative to HU-only groups (*p<0.05) and sham (**p<0.01), although other changes did not significantly differ from sham controls. In contrast, male mice exhibited robust mucin production across multiple exposure conditions, including HU (**p<0.01 JE, ****p<0.0001 IL), HU+50cGy (*p<0.05 JE, **p<0.01 IL), and HU+100cGy (**p<0.01 JE, ***p< 0.001 IL) relative to sham (**Fig. 6E-F**). Notably, mucin levels in HU-exposed males were significantly higher than those observed in HU-exposed females (#p<0.05 JE, ###p<0.001 IL).

Together, these findings indicate that simulated spaceflight conditions disrupt small-intestinal morphology and trigger compensatory mucin production, with particularly strong responses under combined radiation and microgravity exposures. The observed sex-dependent increases in mucin secretion, especially in males, may reflect a protective response to barrier disruption or persistent intestinal inflammation induced by simulated spaceflight stressors.

### 3.6. Simulated spaceflight alters intestinal barrier integrity and promotes leukocyte infiltration in a region- and sex-dependent manner

To further characterize intestinal inflammation associated with simulated spaceflight, we evaluated epithelial barrier integrity and immune cell infiltration by immunofluorescence staining of the tight-junction protein zonula occludens-1 (ZO-1) and the pan-leukocyte marker CD45 in the JE, IL, and colon (**Fig. 7A-I**). ZO-1 expression was used as an indicator of epithelial barrier integrity, whereas CD45 staining quantified immune cell infiltration within the intestinal mucosa. In the JE, both sexes exhibited trends toward decreased ZO-1 expression following simulated spaceflight exposures (**Fig. 7B**). However, these reductions reached statistical significance only in female mice, where ZO-1 levels decreased significantly after 50cGy radiation (****p<0.0001), 100cGy (*p<0.05), and HU+100cGy exposure (*p<0.05) compared with sham controls. In contrast, male mice did not show significant changes in ZO-1 expression in this region compared to sham, yet it was significantly higher than in females after 50cGy (*p<0.05). Immune infiltration in the JE displayed sex-specific patterns (**Fig. 7C**). Male and female mice exhibit increased trends of JE CD45⁺ leukocyte infiltration following HU+IR exposure compared with sham, but no significance. Similar trends were observed in the IL. ZO-1 expression remained largely unchanged in male mice, whereas female mice exhibited a significant decrease after 50cGy radiation (**p<0.01) compared with sham (**Fig. 7E**). In contrast, immune infiltration was markedly elevated in both sexes but through different exposure conditions (**Fig. 7F**). Male mice displayed a significant increase in CD45⁺ cells following 100cGy radiation (*p<0.05) compared with sham and showed significantly greater infiltration than females exposed to the same radiation dose (#p<0.05). Female mice exhibited increased ileal immune infiltration following HU exposure (*p<0.05), with similar trends in the HU+IR groups. In the colon, ZO-1 expression showed significant decreases following 100cGy radiation exposure (*p<0.05), which diminishes in HU+IR groups (**Fig. 7G-H**). In contrast, CD45⁺ immune infiltration was robust in this region (**Fig. 7I**). Female mice showed marked increases in leukocyte infiltration following HU+50cGy exposure (**p<0.01) compared with sham and with males exposed to the same condition (#p<0.05). HU exposure produced significant increases in males relative to HU-exposed females (#p<0.05). Further, 100cGy exposure produced marked increases in males compared to sham (**p<0.01) and to 100cGy-exposed females (###p<0.001). Collectively, these findings demonstrate that simulated spaceflight induces intestinal inflammation characterized by epithelial barrier disruption and immune cell infiltration, with region-specific and sex-dependent responses. Radiation exposure produced more pronounced tight-junction disruption, whereas simulated microgravity and combined HU+IR exposures were associated with stronger leukocyte infiltration, suggesting distinct mechanisms by which spaceflight-related stressors contribute to intestinal immune activation.

**Figure 7.**
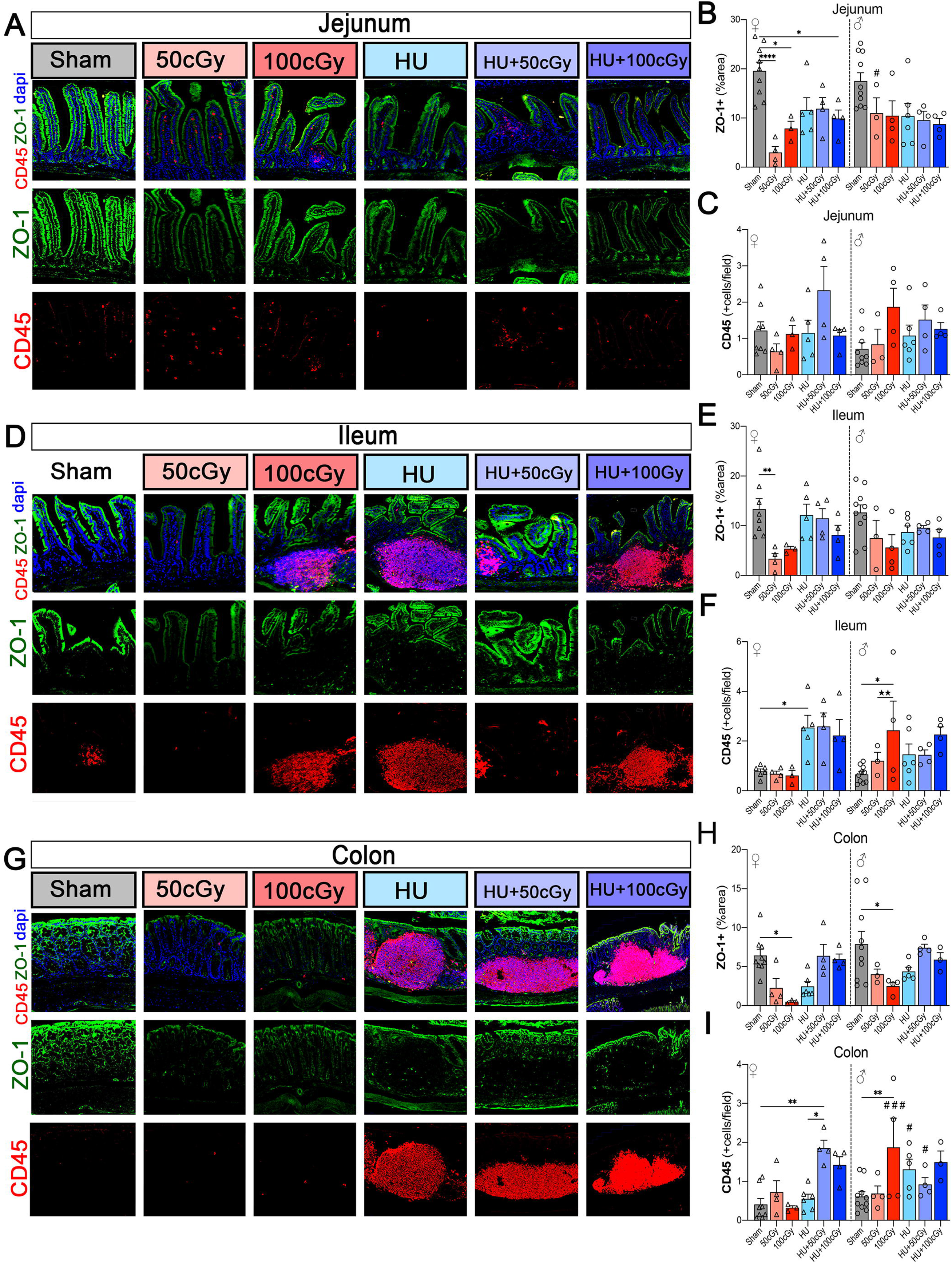
Spaceflight simulation alters intestinal tight junction integrity and leukocyte infiltration in a region- and sex-dependent manner. (A, D, G) Representative immunofluorescence images of the jejunum (A), ileum (D), and colon (G) from mice exposed to Sham, ionizing radiation (IR; 50 or 100cGy), hindlimb unloading (HU), or combined HU+IR (HU+50cGy and HU+100cGy). Sections were stained for ZO-1 (green; tight junction protein), CD45 (red; pan-leukocyte marker), and DAPI (blue; nuclei). Merged images are shown in the top row, with individual ZO-1 and CD45 channels below. (B, E, H) Quantification of ZO-1-positive area (% area) in the jejunum (B), ileum (E), and colon (H). (C, F, I) Quantification of CD45-positive infiltration in the jejunum (C), ileum (F), and colon (I). Data are shown separately for female and male mice. Representative images depict the largest response among males and females. Bars represent mean ± SEM, and symbols indicate individual animals. Statistical significance was determined using two-way ANOVA with appropriate multiple-comparisons testing, with statistical significance indicated: *p<0.05, **p<0.01, ***p<0.001, ****p<0.0001. Statistical significance in sex differences is shown as #p<0.05, ##p<0.01, ###p<0.001. Statistical significance for IR dose effect is shown as l1p<0.05. HU, hindlimb unloading; IR, ionizing radiation; ZO-1, zonula occludens-1.

### 3.7. Simulated spaceflight induces behavioral alterations in affective and cognitive domains

To determine whether simulated spaceflight conditions affect emotional behavior, motor performance, and cognitive function, mice were subjected to a behavioral testing battery following exposure (**Fig. 8**). Anxiety-like behavior was assessed using the elevated plus maze (EPM), in which greater time spent in the closed arms is interpreted as increased anxiety-like behavior. In female mice, time spent in the closed arms showed mild radiation dose-dependent trends, although not statistically significant (**Fig. 8A**). Male mice did not show significant differences across exposure groups, however, the HU+100cGy group was reduced relative to females of the same exposure, indicating a sex-dependent response (##p<0.01). These findings suggest that anxiety-like behavior is sex-specific following SSM. Depressive-like behavior was assessed using the forced swim test (FST), in which increased immobility time reflects greater behavioral despair or passive coping. Male and female mice showed an increased immobility after simulated spaceflight exposures relative to sham, particularly in the HU-exposed group (**p<0.01 females, *p<0.05 males) (**Fig. 8B**). Overall, these findings indicate that simulated spaceflight exposure promotes depressive-like behavior, regardless of sex. Motor coordination and endurance were evaluated using the rotarod test, with performance expressed as the latency to fall at baseline and day 16 (SSM+2) (**Fig. 8C**) and change in latency to fall (individual Δtime from baseline) (**Supp. Fig. 3A**). No significant changes were observed in motor function. However, more sensitive Δtime assessments reveal trends of worsened motor performance following more extreme radiation doses (100cGy) and HU induction (HU, HU+100cGy). In contrast, 50cGy and HU+50cGy exposures display trends of increased motor performance (**Fig. 8C, Supp. Fig 3A**). Overall, these findings display motor adaptability following SSM.

**Figure 8.**
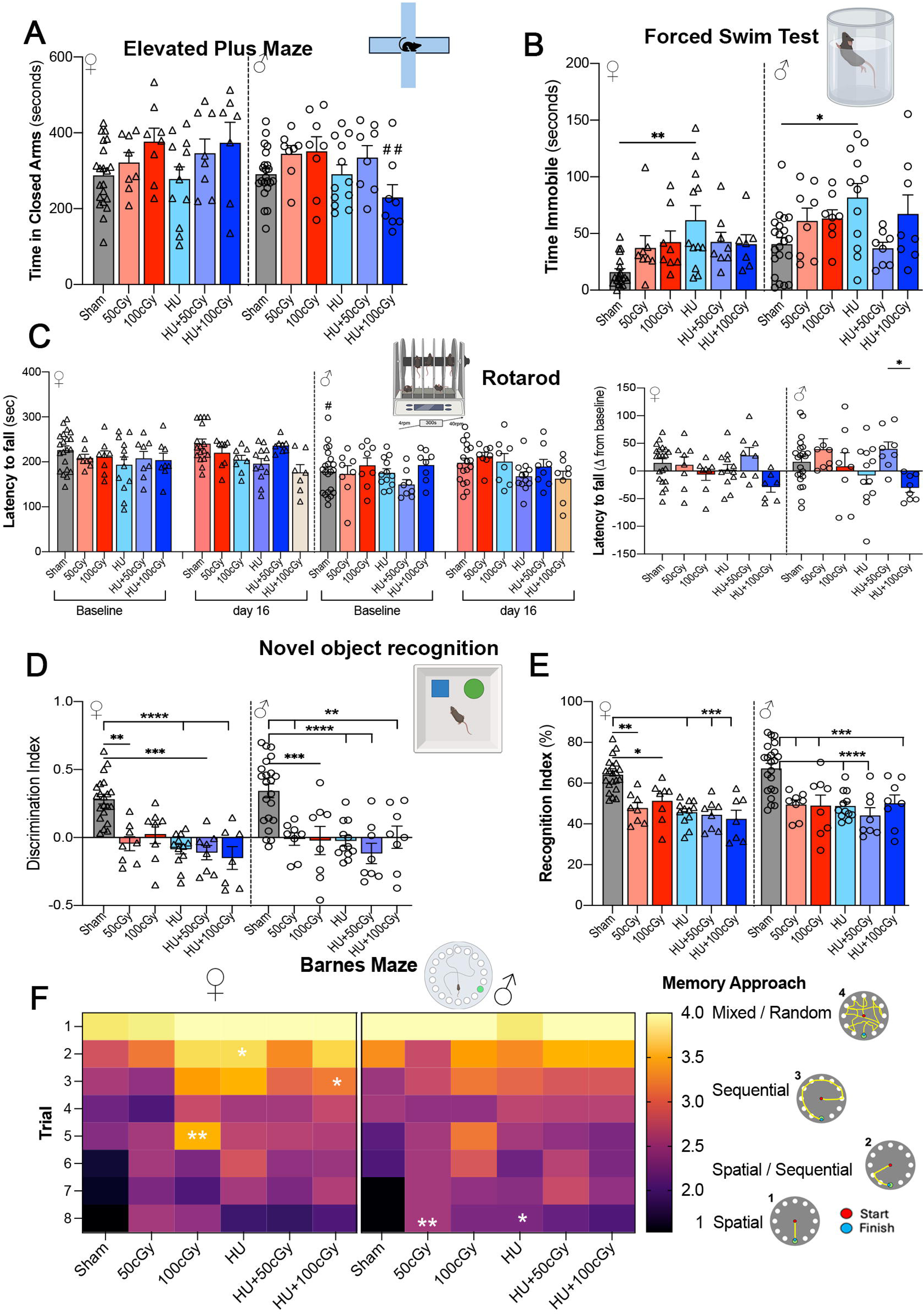
Spaceflight simulation behavioral despair, recognition memory, and spatial search strategies. Mice exposed to Sham, ionizing radiation (IR; 50 or 100cGy), hindlimb unloading (HU), or combined HU+IR (HU+50cGy and HU+100cGy) underwent a behavioral testing battery following the exposure period. (A) Elevated plus maze (EPM): time spent in the closed arms (s), shown separately for female and male mice, as an index of anxiety-like behavior. (B) Forced swim test (FST): immobility time (s), shown separately by sex, as a measure of behavioral despair-like responses. (C) Rotarod: latency to fall at 40 rpm expressed relative to baseline, shown separately for female and male mice, to assess motor coordination and endurance. (D–E) Novel object recognition (NOR): discrimination index (D) and recognition index (%) (E), shown separately by sex, as measures of recognition memory. (F) Barnes maze: heatmap summarizing search strategy across trials in female and male mice, with representative search patterns shown at right. Strategies were categorized from spatial to mixed/random approaches. Bars represent mean ± SEM, and symbols indicate individual animals. Violin plots show data distribution with individual animals overlaid. Statistical significance was determined using one-way ANOVA with appropriate multiple-comparisons testing, with statistical significance indicated: *p<0.05, **p<0.01, ***p<0.001, ****p<0.0001. Statistical significance in sex differences is shown as #p<0.05, ##p<0.01. HU, hindlimb unloading; IR, ionizing radiation; EPM, elevated plus maze; FST, forced swim test; NOR, novel object recognition.

Recognition memory was evaluated using the novel object recognition (NOR) task by calculating the discrimination index and recognition index. In female mice, the discrimination index was significantly reduced across all exposure groups relative to sham, including after 50cGy (**p<0.01), HU (****p<0.0001), HU+50cGy (***p<0.001), and HU+100cGy (****p<0.0001), indicating impaired object discrimination following both single and combined simulated spaceflight stressors (**Fig. 8D**). In male mice, the discrimination index was also significantly reduced relative to sham after 50cGy (**p<0.01), 100cGy (***p<0.001), HU, HU+50cGy (****p<0.0001), and HU+100cGy (**p<0.01). Consistent with these findings, the recognition index in males was significantly lower in the 50cGy, 100cGy, HU+100cGy (***p<0.001), HU, and HU+50cGy (****p<0.0001) groups compared with sham (**Fig. 8E**). Likewise, female mice exposed to 50cGy (**p<01), 100cGy (*p<0.05), HU, HU+50cGy, and HU+100cGy (***p<0.001) had a significantly lower recognition index compared to sham (**Fig. 8E**). Together, these data indicate that simulated spaceflight impairs recognition memory, with particularly pronounced deficits after combined HU and radiation exposure.

Spatial learning and memory were further evaluated using the Barnes maze (BM) by classifying search strategies using a memory approach scoring system (1-4), where lower scores indicate more efficient spatial strategies and higher scores reflect mixed or random search patterns. Across trials, sham mice showed a progressive transition toward spatial strategies, reflected by decreasing scores over time (**Fig. 8F**). In female mice, several exposure groups exhibited altered learning trajectories compared with sham. Radiation exposure (100cGy) produced significantly higher strategy scores during early trials (**p<0.01), indicating reliance on less efficient search strategies, while HU and HU+100cGy also showed elevated scores during intermediate trials (*p<0.05). In male mice, strategy scores were significantly elevated in the 50cGy (**p<0.01) and in the HU group (*p<0.05) during the final trial (trial 8), suggesting delayed adoption of spatial navigation strategies. Mice also exhibit slowed learning through significant reductions in latency to entry (sec) at 50cGy (*p<0.05 males) and 100cGy (*p<0.05 males, ***p<0.001 females) (**Supp. Fig 3C**). At the probe test, mice exhibited no differences in total target time (**Supp. Fig 3B**). Together, these findings indicate that simulated spaceflight exposures disrupt normal spatial learning patterns, producing delayed or inefficient search strategies during BM acquisition.

## 4. DISCUSSION

Spaceflight exposes astronauts to a unique combination of environmental stressors, including microgravity, ionizing radiation, and physiological stress, which together perturb immune, vascular, and neurological systems^8,11,18^. Using a simulated spaceflight model combining HU and IR model, we show that these stressors induce coordinated, multi-organ dysfunction spanning the immune system, gut, BBB, and brain, with associated behavioral impairments Importantly, our data support a systems-level model in which simulated spaceflight disrupts the immune-gut-brain axis, producing sustained inflammatory signaling coupled to barrier failure across peripheral and central compartments (**Fig. 9**).

**Figure 9.**
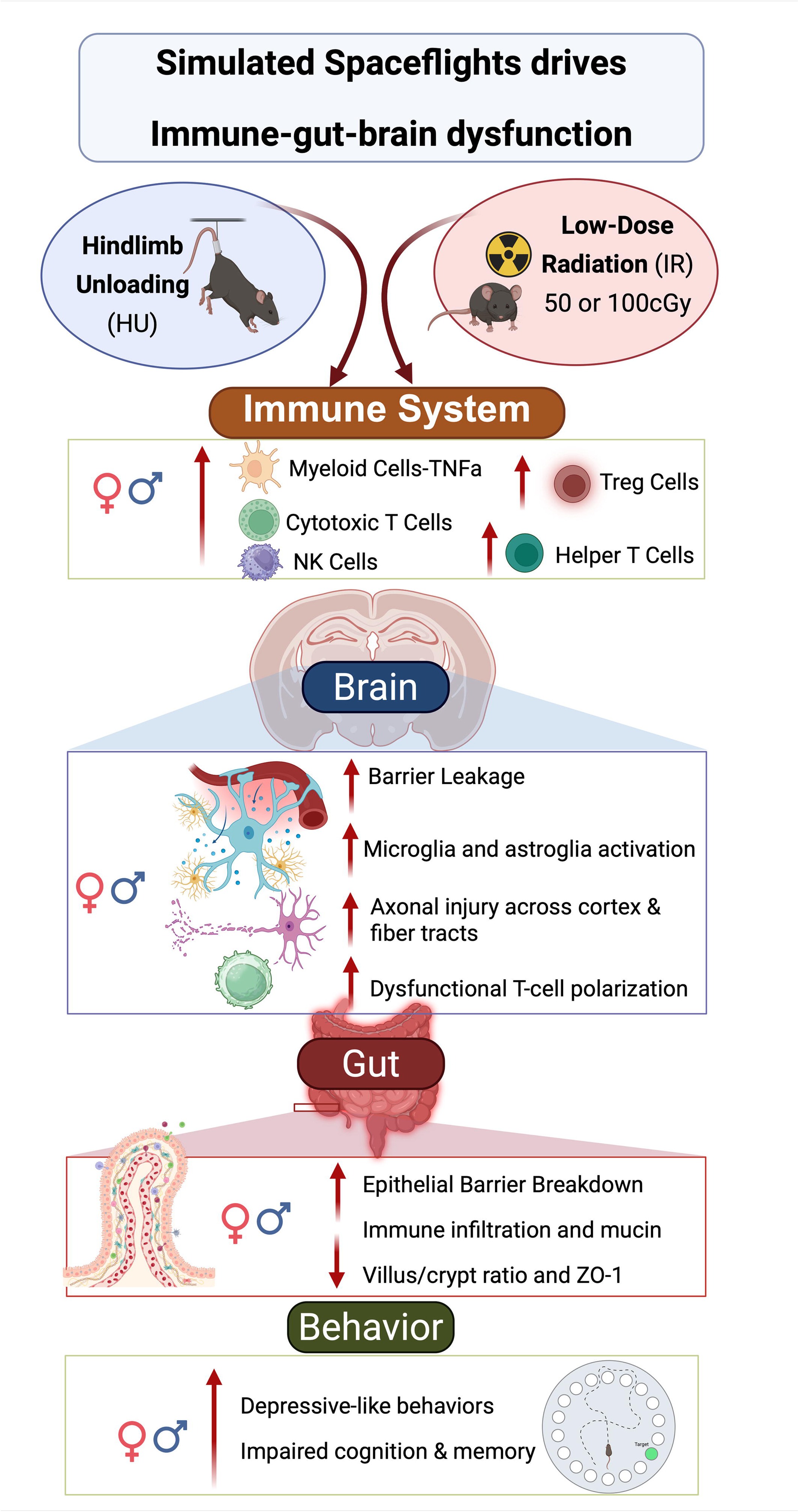
Simulated spaceflight disrupts the immune-gut-brain axis. A combined model of hindlimb unloading (HU) and low-dose ionizing radiation (IR; 50 or 100cGy) induces peripheral immune dysregulation, neuroinflammation, axonal injury, intestinal barrier disruption, and behavioral deficits. Alterations include increased inflammatory and T-cell responses, blood-brain barrier leakage, microglial and astroglial activation, epithelial breakdown, reduced villus/crypt ratio and ZO-1, and anxiety-like, depressive-like, cognitive, and motor impairments.

Systemically, simulated spaceflight induced a rapid and sustained immune imbalance characterized by early innate activation followed by persistent adaptive dysregulation^19,21^. Radiation acted as a primary trigger of acute myeloid activation, while combined HU+IR exposure prolonged immune perturbations, particularly within T cell and NK cell compartments. The delayed NK cell activation observed under combined conditions suggests that microgravity does not simply add to radiation effects, but modifies immune set points, potentially by altering trafficking, activation thresholds, or cytokine responsiveness. Mechanistically, this pattern is consistent with a transition from an acute inflammatory response to a maladaptive, unresolved immune state resembling features of T cell exhaustion^19,21,53^. Thus, simulated spaceflight appears to shift immune responses from transient activation toward chronic, low-grade inflammatory persistence, a state known to impair tissue repair and promote neurodegeneration.

A key determinant of this response was biological sex^54^. Females exhibited greater inflammatory activation and more pronounced barrier leakage in both brain (IgG extravasation) and gut (CD45⁺ infiltration), indicating heightened systemic and neurovascular sensitivity to spaceflight stressors. These findings suggest that females may mount a more rapid or amplified immune response, but at the cost of increased barrier vulnerability. Rather than being purely protective, this heightened responsiveness may predispose to immune-driven tissue damage when regulatory resolution fails. This interpretation is consistent with prior spaceflight and GCR simulation studies and supports the concept that sex-dependent neuroendocrine-immune coupling critically shapes susceptibility to spaceflight-induced pathology^20,54,55^. These results underscore that sex is not simply a modifier, but a determinant of pathophysiological trajectory under spaceflight conditions.

At the level of the CNS, neuronal structural integrity was preferentially impacted by radiation exposure. Increased axonal phosphorylation (SMI-312) and reduced dendritic complexity (MAP-2) indicate that radiation disrupts both axonal maintenance and dendritic stability. Radiation-induced metabolic stress may therefore shift neurons toward survival signaling pathways that alter axonal maintenance and repair^56^. These changes likely reflect metabolic stress-induced shifts in neuronal homeostasis, in which survival signaling pathways are engaged at the expense of effective repair and proteostasis. Spatial transcriptomic mapping of mice flown on the ISS supports our cortical and white matter track findings, with exacerbated neurodegenerative pathways involving impacted synaptic transmission and protein misfolding^57^. Similar metabolic alterations have been reported in spaceflight models, including increased glycolysis and altered lipid metabolism in neural and immune cells^19,58^. This imbalance may transiently preserve neuronal structure while promoting the accumulation of dysfunctional axonal elements, ultimately impairing network function. The preferential vulnerability of white matter tracts, including the CC, further suggests that metabolically demanding, oligodendrocyte-supported regions are particularly sensitive to spaceflight stressors.

Our results show a clear sex-dependent neurotrophic response to simulated spaceflight. Cortical BDNF levels increased in males under HU+100cGy, whereas no changes were observed in females. This suggests that males engage in a compensatory neuroprotective response^59^ to counteract inflammatory and neuronal stress^60^, while females may lack this adaptive mechanism, potentially contributing to their greater vulnerability to barrier dysfunction and neuroinflammation. Previous spaceflight studies report no significant changes in BDNF^61^, whereas ground-based HU models show reductions^62^, indicating impaired neurotrophic support under microgravity. Together, these findings suggest that combined microgravity and radiation induce a sex-dependent modulation of neurotrophic signaling, rather than a uniform effect on BDNF.

Converging with these structural alterations, we observed robust and region-specific neuroinflammation. In line with prior findings, mice flown on the International Space Station (ISS) display pronounced metabolic dysfunction, neuronal injury, apoptosis, and oxidative stress signatures within the hippocampus^17^. In our model, microglial activation and astrocyte reactivity were most prominent following combined HU and radiation exposure, particularly within the CA1 and DG subregions, key areas for memory and cognitive processing that are highly sensitive to inflammatory stress. Supporting these observations, longitudinal studies of simulated GCR exposure exceeding 100 days demonstrate sustained elevations in Iba-1 expression in the hippocampus^63^, indicative of persistent neuroinflammation. Such chronic activation, observed both at subacute (e.g., 14 days) and extended time points, suggests impaired immunoregulatory resolution and raises concern for long-term cognitive dysfunction and neurodegenerative risk. Mechanistically, radiation-induced metabolic stress may shift neurons toward survival-oriented signaling pathways, potentially altering axonal maintenance and repair processes^64^. Even low-dose radiation may exert similar cumulative effects, further exacerbating neuronal vulnerability. Notably, white-matter regions such as the CC also exhibited robust microglial activation, indicating that interhemispheric projection pathways may be particularly susceptible to simulated spaceflight stressors^57^. Spatial transcriptomic analyses further reveal that astrocyte-driven neuroinflammatory responses are compounded by reduced accessibility of immunoregulatory transcription factor motifs^57^, suggesting epigenetic constraints on the resolution of inflammation. Together, these findings indicate that neuronal structural injury and glial activation occur concurrently and may reinforce one another during prolonged exposure to spaceflight-like conditions.

BBB dysfunction emerged as a central interface linking systemic inflammation to CNS pathology^21,65,66^. Sex-dependent differences were also evident at the BBB level, with females exhibiting greater leakage, consistent with their heightened inflammatory profile. Structural disruption, indicated by reduced claudin-5, was primarily radiation-driven, whereas functional leakage (IgG extravasation) occurred even in regions without overt tight junction loss. This dissociation indicates that BBB permeability is not solely dependent on tight junction integrity but also involves dynamic regulatory processes, including endothelial transport, oxidative damage, and lipid membrane instability. Consistent with this, mice exposed to spaceflight conditions on the ISS show molecular markers of BBB dysfunction, including increased PECAM-1 and AQP-4 and reduced ZO-1^4,17^. Notably, HU alone induced substantial IgG leakage, suggesting that microgravity-related hemodynamic and vascular alterations are sufficient to compromise BBB function independently of structural breakdown^49^. This supports a model in which BBB disruption arises from both structural injury and functional dysregulation, with distinct contributions from radiation and microgravity. HU exposure frequently produced pronounced IgG leakage into the brain parenchyma, even when tight-junction loss was less apparent. This observation suggests that BBB permeability may alternatively be influenced by microgravity-related vascular or metabolic factors^49^. Cephalad fluid shifts alter vascular pressure, oxidative stress, and endothelial transport changes have all been proposed as contributors to BBB leakage under microgravity conditions. Importantly, females frequently exhibited greater IgG leakage than males, suggesting increased neurovascular vulnerability.

Heat shock proteins are known to stabilize damaged proteins and buffer cellular stress, particularly under microgravity-like conditions^67,68^. HSP90 increased in male mice after radiation, while HSP25 was elevated in females only under combined HU+100cGy, indicating activation of stress-response pathways to protect BBB integrity^69^. Despite this, these responses were insufficient to prevent neuroinflammation and BBB disruption. In females, HSP90 and HSP25 were reduced under HU conditions, suggesting a loss of baseline protective function rather than a beneficial effect. This downregulation likely reflects sustained cellular stress^70^ and contributes to BBB breakdown^71^. These results highlight sex-dependent differences in stress-response pathways underlying neuroinflammation and BBB dysfunction during simulated spaceflight.

Parallel disruption of the intestinal barrier reinforces the central role of the GBA in mediating systemic and CNS responses. Reduced villus-to-crypt ratios and decreased ZO-1 expression indicate structural and functional epithelial compromise, while increased CD45⁺ infiltration reflects active intestinal inflammation. Interestingly, radiation exposure produced stronger tight-junction disruption, whereas HU and combined exposures were associated with greater immune infiltration. Importantly, different stressors produced distinct patterns: radiation preferentially disrupted tight junction integrity, whereas microgravity promoted immune cell infiltration, suggesting that microgravity-like conditions impose substantial stress on intestinal architecture^40^. This suggests that spaceflight stressors converge on the gut through partially independent but complementary mechanisms, together amplifying barrier dysfunction. In addition, mucin production increased in a sex-dependent manner, particularly in females under combined exposures. While mucin secretion provides a protective barrier against luminal pathogens, excessive production may reflect unresolved mucosal inflammation. Such inflammatory states can impair microbial function and digestive processes^72^. Although intestinal architecture remains underexplored in spaceflight studies, prior reports have documented disruptions in metabolism, neuroendocrine signaling, and gene expression^73,74^, further supporting the vulnerability of the gut under spaceflight-like conditions. Given the rapid bidirectional communication between the gut and brain, intestinal permeability is likely an upstream driver of systemic immune activation and subsequent neuroinflammation.

These multi-system alterations translate into measurable behavioral deficits. Cognitive impairments, particularly in recognition memory, align closely with hippocampal inflammation, BBB disruption, and neuronal injury observed in our model. Simulated deep-space exposure is widely supported in the literature as a driver of deficits in learned helplessness, memory, cognitive accuracy, and processing spee^75–79^. The absence of strong anxiety-like changes, coupled with increased depressive-like behavior, suggests selective vulnerability of mood-related circuits. Likewise, longitudinal NOR measurements (80 days) in mice exposed to lunar (15 cGy) and Martian (50 cGy) equivalent simulated GCR show diminished recognition in males but not in females. Notably, the spatial distribution of radiation-induced reactive species differs by radiation type, leading to distinct behavioral outcomes; low-LET radiation has been associated with greater behavioral impairment in some contexts^80^.

Importantly, our data indicate that radiation and microgravity exert distinct but interacting effects. Radiation predominantly drives neuronal injury and tight junction disruption, while microgravity contributes to vascular leakage, immune infiltration, and intestinal dysfunction. When combined, these stressors do not act additively but rather synergistically amplify systemic and neuroinflammatory responses, creating a persistent pathological state characterized by impaired resolution and multi-organ dysfunction.

These findings have direct implications for astronaut health during long-duration missions. The identification of immune priming by microgravity suggests that early exposure may sensitize individuals to subsequent radiation damage, highlighting the importance of temporal dynamics in risk assessment. Persistent alterations in adaptive immunity further indicate incomplete recovery, raising concerns about long-term immune competence. Several limitations should be considered. The use of low-LET X-ray radiation does not fully recapitulate the high-LET particle environment of galactic cosmic radiation, which induces more complex DNA damage^80^. Although dose adjustments approximate biological effectiveness, differences in energy deposition patterns may influence outcomes^1,81^. Additionally, while our model captures key features of spaceflight stress, it does not fully capture the complexity of space conditions.

In summary, our study provides an integrated framework linking immune dysregulation, barrier dysfunction, and neuroinflammation under simulated spaceflight conditions. We demonstrate that combined microgravity and radiation exposure drive a persistent, system-wide inflammatory state that disrupts both peripheral and central homeostasis. These findings identify the immune-gut-brain axis as a critical mediator of spaceflight-induced pathology and highlight potential therapeutic targets, including modulation of immune responses, preservation of barrier integrity, and enhancement of neurovascular resilience.

## Acknowledgements.

The authors thank Juan Fernandez and the HMRI Machine Shop Core for their assistance in constructing the HU cages. Figure 1a and Figure 9 were created with BioRender.com

## Abbreviations

BCA: Bicinchoninic acid
BBB: Blood-brain barrier
BM: Barnes-Maze
CC: Corpus callosum
CG: Cingulate gyrus
cGY: Centigray
CNS: Central nervous system
DG: Dentate gyrus
EC: External capsule
EPM: Elevated plus maze
FST: Forced swim test
GALT: gut-associated lymphoid tissue
GBA: Gut-brain axis
GCR: Galactic cosmic rays
H&E: Hematoxylin and eosin
HU: Hindlimb unloading
IgG: Immunoglobulin G
IL: Ileum
IR: Ionizing radiation
JE: Jejunum
LET: Linear energy transfer
NOR: Novel object recognition
PBST: PBS-Triton X-100
PFA: Paraformaldehyde
PVDF: Polyvinylidene difluoride
SEM: Standard error of the mean
SSC: Somatosensory cortex
SSM: Spaceflight simulation model
ZO-1: Zonula occludens-1

---

**Supplementary Figure 1. Flow cytometry gating strategy for peripheral immune cell populations.** (A) Initial gating strategy for identification of viable single leukocytes. Cells were first selected based on forward scatter (FSC-A) and side scatter (SSC-A) to exclude debris, followed by doublet discrimination using FSC-A versus FSC-H. Viable leukocytes were then identified by gating on CD45⁺ cells. (B) From the CD45⁺ population, innate immune cell subsets were identified. Myeloid cells were gated based on CD11b⁺ expression, and activated myeloid cells were identified using activation markers (TNF-α). Natural killer (NK) cells were identified as NK1.1⁺ cells within the CD45⁺ population and further subdivided into pro-inflammatory NK cells and regulatory NK cells based on TNF-α/IFN-γ or IL-10 cytokine expression, respectively. (C) Adaptive immune cell populations were identified within the CD3⁺ T cell compartment. CD8⁺ cytotoxic T cells and CD4⁺ helper T cells were first separated, after which CD4⁺ cells were further subdivided into Th1, Th2, and regulatory T cells (Treg) based on lineage-defining markers (IFN- γ and IL-4) and transcription factors (FoxP3). Representative pseudocolor plots illustrate the sequential gating strategy used to quantify immune cell populations analyzed in the study.

**Supplemental Figure 2. Blood-brain barrier leakage analysis, sex specific responses, and vessel IgG infiltration.** (A) Blood-brain barrier (BBB) leakage assessment pipeline using QuPath (v0.6.0, x64). An automated vessel classifier script was generated to detect vessels and perivascular shells for IgG detection in those respective areas. (B-E) Sex specific responses from Fig. 5 are depicted for the cingulate gyrus and somatosensory cortex after HU (B,C), in the dentate gyrus after 50cGy (D), and in the CA1 after 100cGy (E). IgG infiltration is depicted through %area IgG expression within defined vessels for the dentate gyrus (F), CA1 (G), cingulate gyrus (H), and somatosensory cortex (I). Data are shown separately for female and male mice. Values represent mean ± SEM, with each dot representing an individual animal. Statistical significance was determined using two-way ANOVA followed by multiple-comparisons testing, with statistical significance indicated: *p<0.05, **p<0.01, ***p<0.001, ****p<0.0001. HU, hindlimb unloading; IR, ionizing radiation; Cldn-5, Claudin-5.

**Supplemental Figure 3. SSM does not alter individual motor performance or time-based spatial memory at the probe test.** (A) Rotarod: latency to fall at 40 rpm expressed as difference from individual mouse baseline (Δtime). (B) Barnes maze probe test performed on day 5, measured as total time at target (sec). (C) Barnes maze training tests (trials) measured through latency to escape box entry. Bars represent mean ± SEM, and symbols indicate individual animals. Violin plots show data distribution with individual animals overlaid. Statistical significance was determined using one-way ANOVA with appropriate multiple-comparisons testing, with statistical significance indicated: *p<0.05, **p<0.01, ***p<0.001. Statistical significance in sex differences is shown as #p<0.05. Statistical significance for IR dose effect is shown as l1p<0.05, l1l1p<0.01. HU, hindlimb unloading; IR, ionizing radiation; EPM, elevated plus maze; FST, forced swim test; NOR, novel object recognition.

