## Supplementary figures and images for "Simulated spaceflight disrupts the immune-gut-brain axis and drives sex-dependent neuroinflammation, axonal injury, and behavioral deficits"

### Supplementary Figure 1

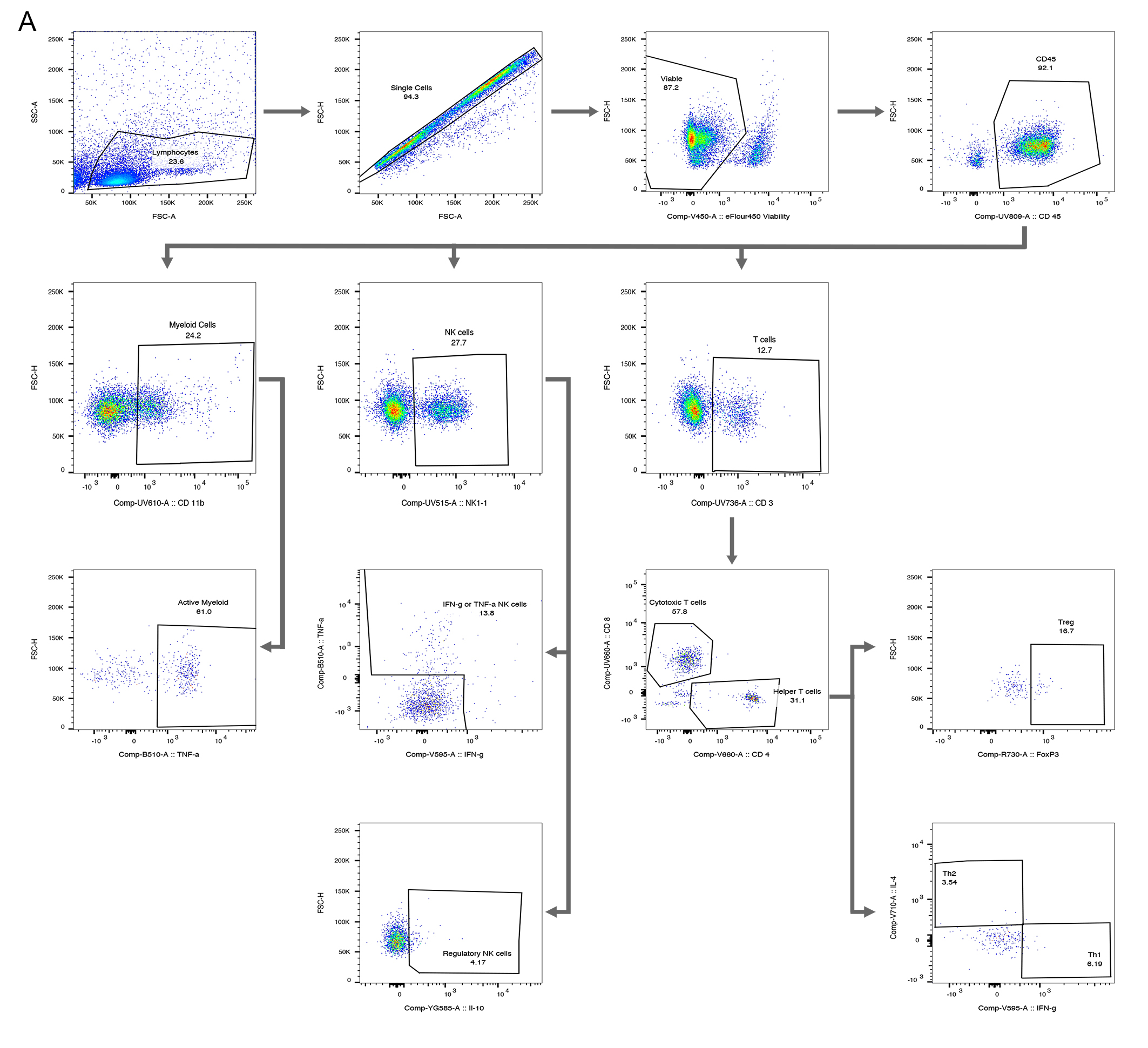

### Supplementary Figure 2

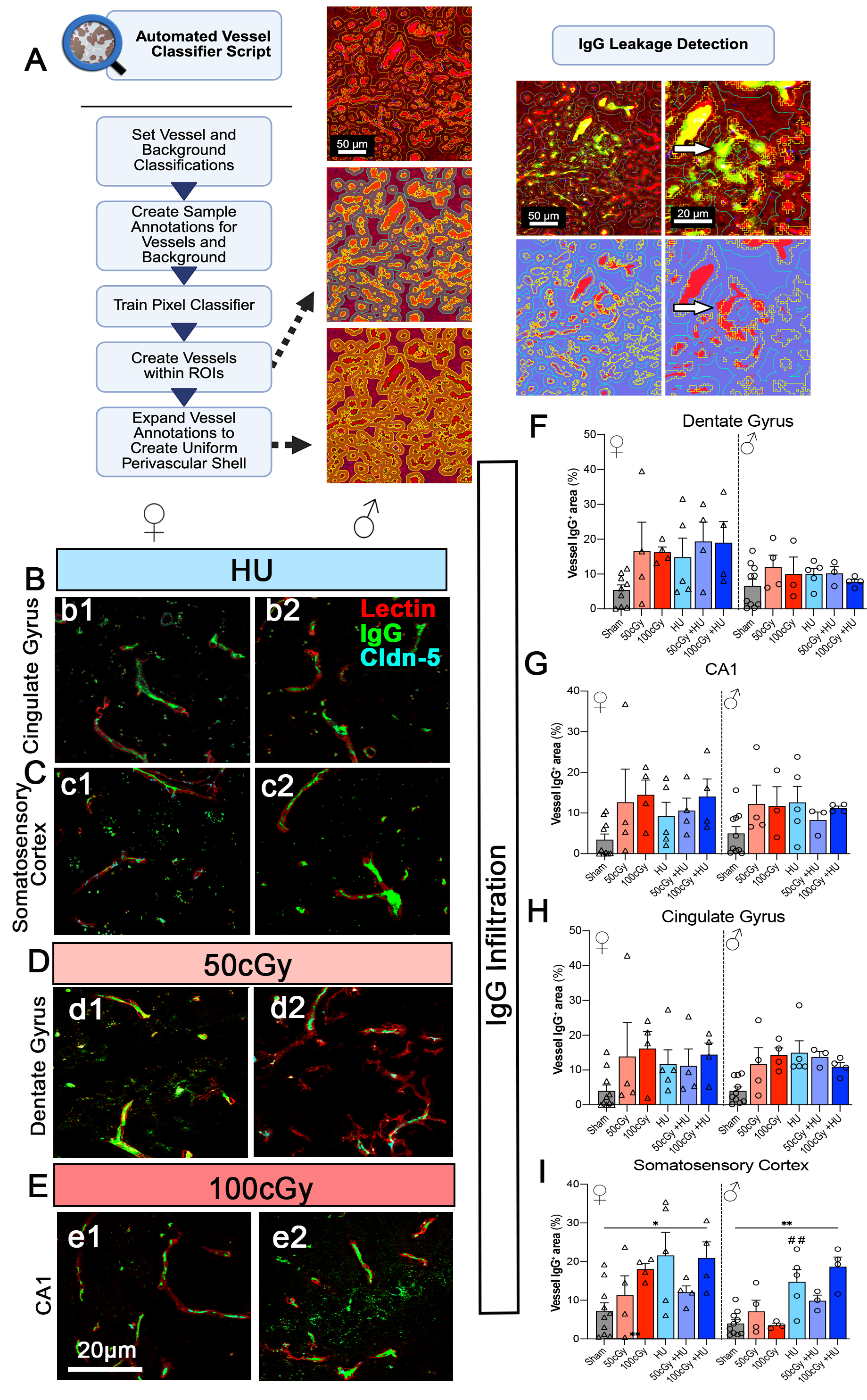

### Supplementary Figure 3

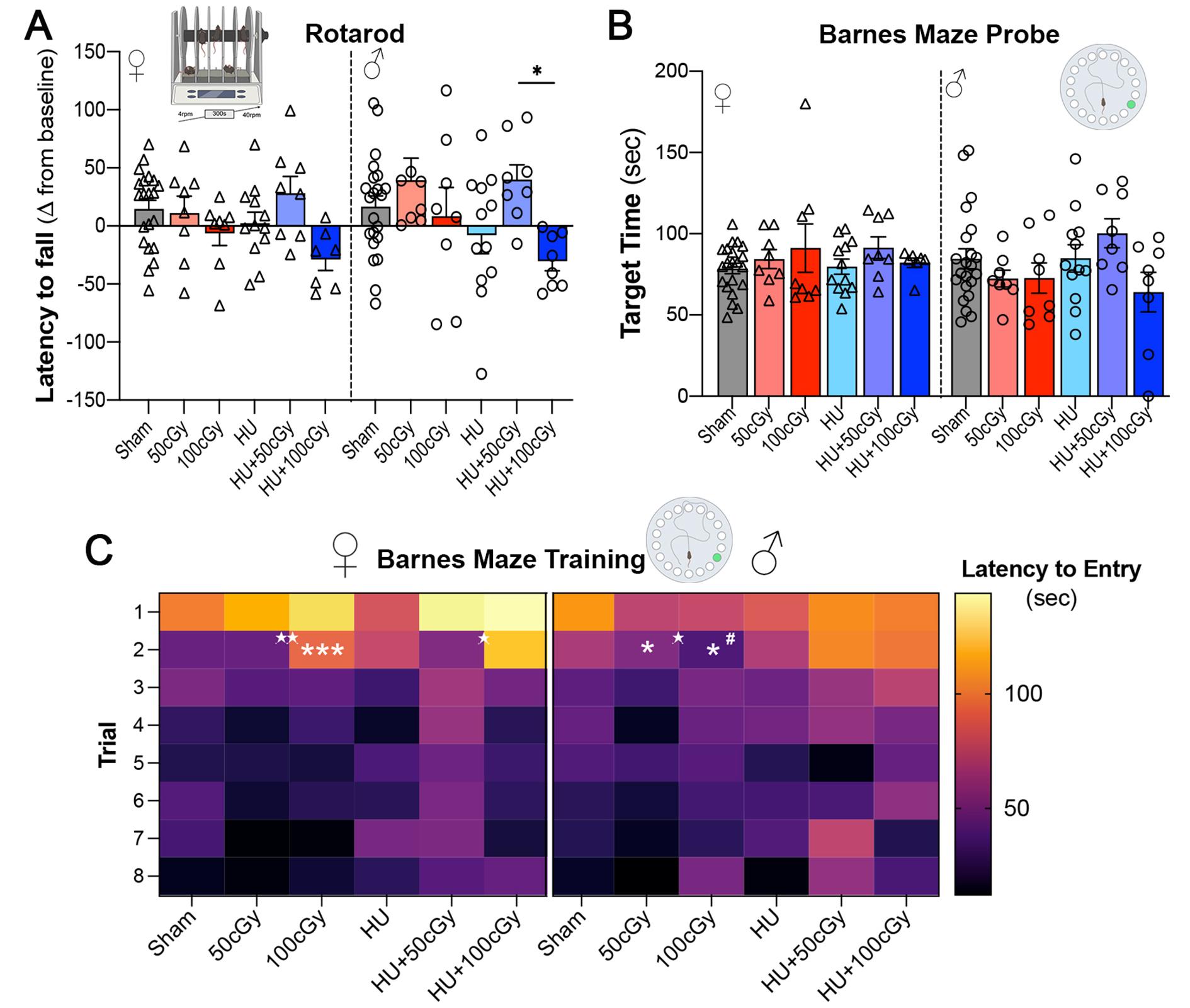
